# A Chemical Mechanistic Path Leads the Way to Cellular Argpyrimidine

**DOI:** 10.1101/2025.05.30.657038

**Authors:** Vo Tri Tin Pham, Suprama Datta, Amy C. Sterling, Stefan M. Hansel, Rebecca A. Scheck

## Abstract

Argpyrimidine (APY) is a methylglyoxal-derived advanced glycation end-product (AGE) that has been associated with multiple diseases. As APY forms without an enzyme, it remains exceptionally difficult to pinpoint where APY is likely to be found, both on individual proteins and in cells. In this study, we used a peptide model system and mass spectrometry analysis to investigate the chemical mechanism through which APY arises from methylglyoxal (MGO), a biologically relevant glycating agent. Consistent with other proposed APY formation mechanisms, our results identify AGE species with a mass change of [M+144], presumably including tetrahydropyrimidine (THP), as a direct precursor to APY. However, our results rule out previously proposed reductone or oxidative decarboxylation mechanisms. Instead, we show that a formal oxidation step is not required, and that formate is released instead of CO_2_. We further show the potential for a nearby residue such as Tyr to assist in the APY formation mechanism by acting as a general base. These experiments also reveal that phosphorylated Tyr or Ser residues can also promote equivalent levels of APY formation, despite introducing additional negative charges that we previously showed to impede glycation. Guided by these mechanistic insights and a newly defined role for phosphorylated residues on glycation substrates, we performed quantitative bottom-up proteomics analysis for MGO-treated cells. Gene ontology analysis for AGE-modified proteins revealed enrichment of phosphorylation-related terms (e.g. kinase activity or protein phosphorylation) for APY, while other Arg post-translational modifications did not. Collectively, these data define a chemical mechanistic path to APY and suggest significant crosstalk between cellular phosphorylation and glycation events including APY formation.

## Introduction

Argpyrimidine (APY) is one of a group of nearly 40 protein post-translational modifications (PTMs) known as advanced glycation end-products (AGEs).^1–3^ AGEs are hallmarks of aging and disease that form non-enzymatically on Lys, the N-terminus, or—in the case of APY—on Arg residues.^1–3^ Despite a relatively high p*K*_a_, the Arg guanidino group reacts selectively with ⍺-oxoaldehydes such as the biological glycating agent methylglyoxal (MGO), forming multiple AGEs.^4–6^ In addition to APY, these include the methylglyoxal-derived hydroimidazolone isomers (MGH-1, -2, & -3), dihydroxyimidazolidine (MGH-DH), carboxyethylarginine (CEA), and tetrahydropyrimidine (THP) (**Figure 1**). Although there is accumulating evidence that certain AGEs, especially the MGH isomers and CEA, lead to specific biological consequences,^7–12^ far less is known about the biology of APY. Several reports have correlated APY with cellular processes like apoptosis, autophagy, or chaperone activity, and a few have shown its association with diseases like brunescent cataractous lens, filial amyloidotic polyneuropathy, or cancer.^13–21^ Still, there are virtually no causal connections that link APY to specific biological consequences.

**Figure 1.**
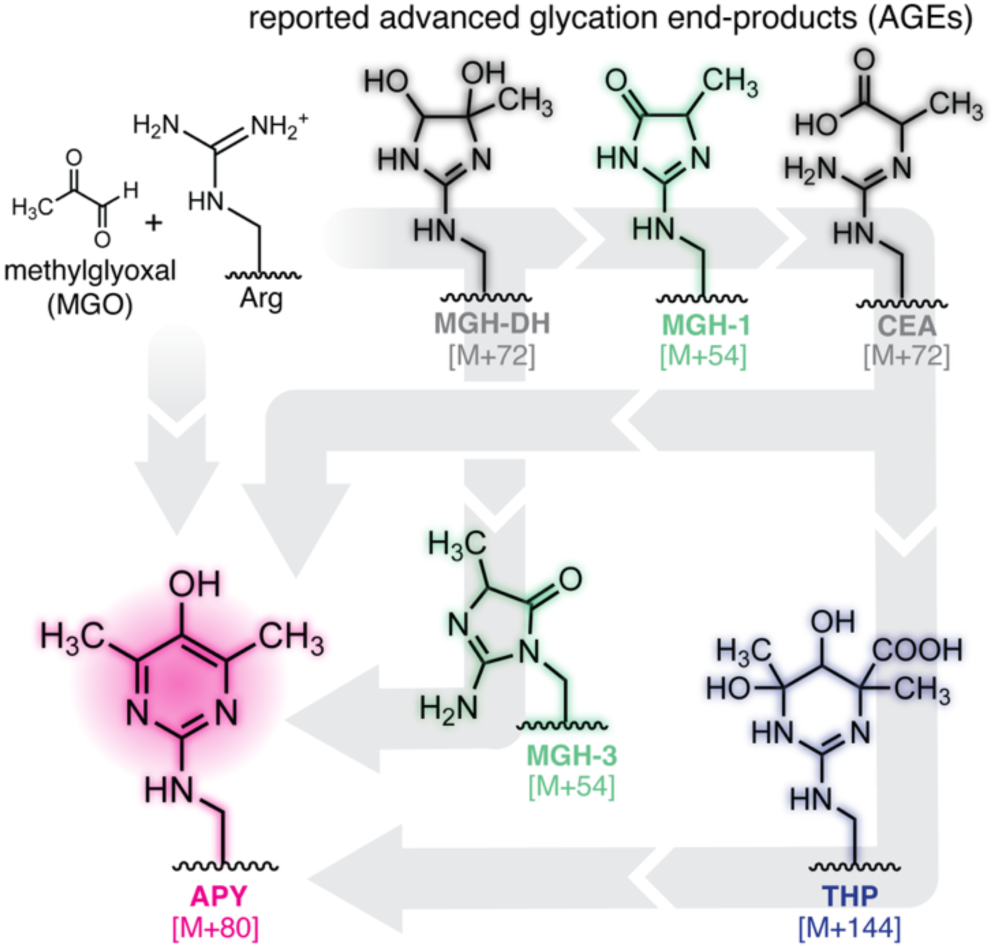
Overview of MGO-derived advanced glycation end-products. In addition to APY, these include the methylglyoxal-derived hydroimidazolone isomers (MGH-1, -2, & -3), dihydroxyimidazolidine (MGH-DH), carboxyethylarginine (CEA), and tetrahydropyrimidine (THP). Several possible pathways for APY formation have been previously proposed (gray arrows).

These open questions persist because APY has been notoriously difficult to track due to a lack of highly specific tools and an unknown mechanism of formation. Given that APY requires more than a single MGO molecule to form, it could be particularly relevant under hyperglycemic conditions. Additionally, it has been proposed that APY formation also depends on oxidative stress, suggesting that it could be a critical link between glycation and other cellular stresses.^20,22,23^ It is therefore essential to understand APY’s formation mechanism to pinpoint the features that influence its formation and the locations—both on proteins and in cells—where APY is more likely to accumulate.

Previous work in our lab has shown that specific Arg-derived AGEs depend not only on MGO concentrations, but also the surrounding chemical microenvironment.^4,5^ We have shown that several of these Arg-derived AGEs are mechanistically connected and can interconvert.^4^ In this work, we use a peptide model system to determine chemical features that favor APY formation. We show that although APY is generated more slowly than other AGEs, it is also quite stable once formed. We also confirm that [M+144] species, presumably including tetrahydropyrimidine (THP), are a direct precursor to APY, and we propose a new APY formation mechanism that involves an initial retro-aldol step, ultimately generating formate as a byproduct, rather than an oxidative decarboxylation. We further demonstrate that the presence of a nearby Tyr can facilitate APY formation. These experiments also revealed that phosphorylated Tyr or Ser residues could also facilitate APY formation, despite adding substantial negative charges that we have previously shown to inhibit glycation.^3–5^ These results prompted us to conduct quantitative proteomics experiments investigating APY formation in live cells treated with MGO, enabling us to draw a previously unknown connection between AGE formation and phosphorylation in a cellular context. These findings provide essential mechanistic insights that are greatly needed for future studies that seek to develop chemical or fluorescent tools that can be used to predict, sense, and study glycation events in living cells.

## Results

### Determining the APY precursor

Although APY requires two molecules of MGO (72.02 Da), its mass change ([M+80]) does not clearly match with an obvious mechanism involving loss or gain of water, as is the case for other AGEs like MGHs, MGH-DH, or CEA.^3^ This has made it particularly challenging to define the APY formation mechanism.^2,24,25^ Prior work from our lab and others’ points to tetrahydropyrimidine (THP, [M+144]) as a possible precursor to APY.^4,24,25^ Specifically, we found that by observing AGE distributions for up to 4 weeks of incubation with MGO, there was an inversely proportional relationship between APY and [M+144] species over time.^4^ We therefore initiated this study with the same hit sequence from our prior study (Ac-LESRHYA, peptide **1**) that was selected from a one-bead one-compound peptide library for its ability to form MGH-1.^4^ We treated peptide **1** (1 mM) with two equivalents (2 mM) of MGO, expecting that APY levels would increase at higher MGO concentrations. However, little to no APY was observed (< 3% APY), even after 72 hours of treatment, despite total glycation nearing 100% after 48 h, as assessed by liquid chromatography-mass spectrometry (LC-MS) (**Figure 2A,B**). In addition to peptide **1**, we also evaluated two additional peptide sequences, both from lens crystallin, that we previously identified as APY-modified using MGO-treated proteins *in vitro* (**Figure S1**).^5^ Even when increasing MGO concentrations to 3 mM, there was little to no (< 3%) APY after 48 hours across all three peptides, signifying that APY formation is largely independent of initial MGO concentrations, despite requiring two MGO equivalents to form.

**Figure 2.**
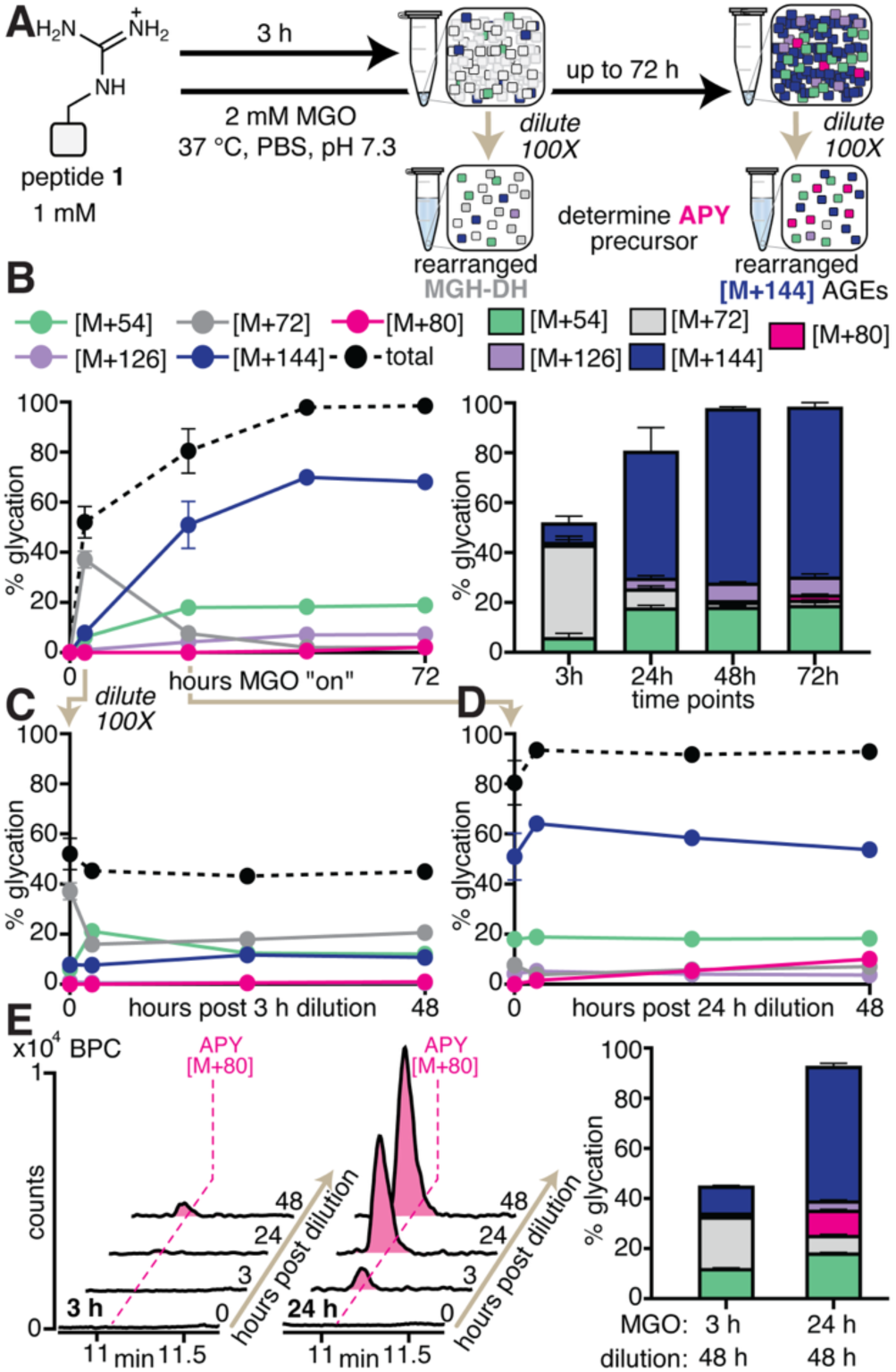
[M+144] AGEs are a Precursor to APY. **(A)** General scheme showing the experimental setup used to determine the APY precursor. **(B)** Peptide **1** (1 mM) was treated with 2 mM MGO and incubated for up to 72 h (labeled as “hours MGO “on”), as shown in a time course (left) and stacked bar graph (right) showing AGE distributions. **(C)** After 3 h, (when MGH-DH became the predominant adduct) or **(D)** after 24 h (when [M+144] species become the predominant AGEs), the reaction was diluted 100X in the same buffer and incubated for additional 48 h post dilution (“hours post dilution”). Since dilution markedly slows MGO addition, this protocol diminishes new AGE formation on already glycated peptides, enabling the clear observation of rearrangements between AGEs. **(E)** Representative base peak chromatograms (BPC) (left) and bar graph (right) for peptide **1** at several post dilution time points (n≥3 for all reactions). Please note that not all error bars are visible, as many are smaller than the size of the symbol. A significant increase in APY [M+80] (magenta) levels was observed at 48 h post dilution of the 24 h time point when compared to that of the 3 h timepoint. These results suggest that [M+144] species (blue), and not MGH-DH (gray), is a precursor for APY.

Nonetheless, the major difference in AGE distribution at 3 and 24 h allowed us to test the hypothesis that [M+144] AGEs (likely including THP) are a precursor to APY. To do so, we incubated peptide **1** with 2 mM MGO for either 3 h or 24 h. After this initial incubation, we diluted the reaction mixture 100X in the same buffer and compared the resulting AGE distributions at several post-dilution time points (**Figure 2C—E**). Since dilution markedly slows MGO addition, this protocol diminishes new AGE formation on already glycated peptides, thereby enabling the clear observation of rearrangements between AGEs. At 3 h, MGH-DH (37.1 ± 3.4 %) was the predominant AGE, but at 24 h, multiple [M+144] species (51.0 ± 9.4 %) became the predominant AGEs. While dilution of the 3 h incubation did not yield any APY even at 48 h post dilution (**Figure 2C,E**), we observed a substantial increase in [M+80] levels when the 24 h incubation was diluted, reaching 10.0 ± 0.3% APY after 48 h post dilution (**Figure 2D,E**). These data confirmed the hypothesis that [M+144] AGEs are a precursor to APY. We note that we consistently observed up to four different [M+144] species with distinct retention times (**Figure 3 & Figure S2**). To our knowledge, THP is the only [M+144] AGE that has been reported.^26^ While it is possible that THP diastereomers may have different retention times, we also suspect that these discrete [M+144] adducts could also represent other intermediate structures or other double addition adducts (see also **Figure 4**).

**Figure 3.**
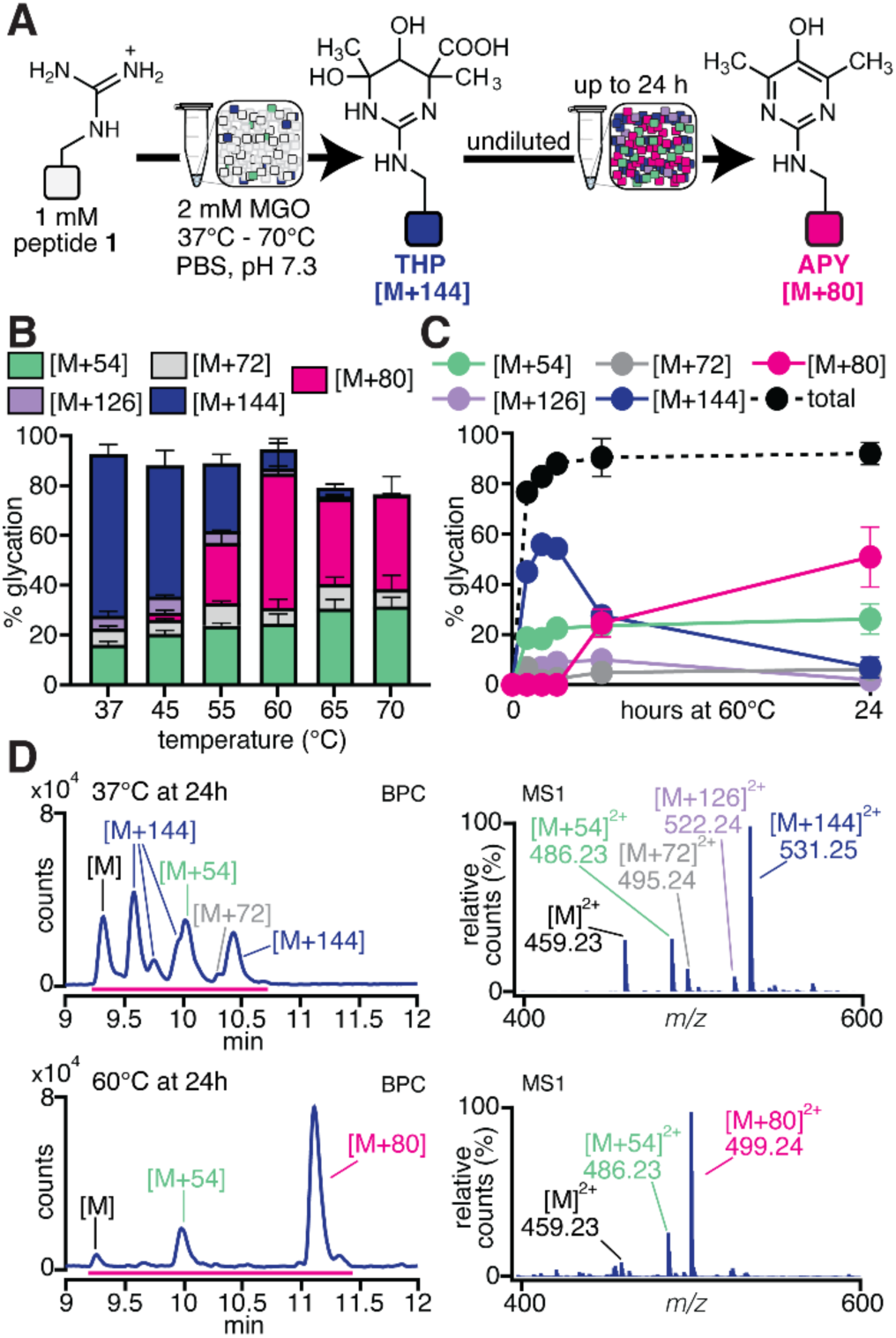
APY is a stable AGE. **(A)** Seeking to increase APY levels, we conducted a temperature scan by treating peptide **1** with 2 mM MGO over a range of temperatures from 37 °C to 60 °C. The reaction was incubated for 24 h undiluted, and AGE distributions were assessed *via* liquid chromatography-mass spectrometry (LC-MS) analysis. **(B)** Stacked bar graph showing AGE distributions for peptide **1** treated with MGO for 24 h at 37 °C, 45 °C, 55 °C, 60 °C, 65 °C, and 70 °C. At 55 °C, we began to observe a notable increase in APY [M+80] (magenta) that was concomitant with reduced [M+144] (blue) levels. APY was the predominant adduct in just 24 h at 60 °C. **(C)** Time course of peptide **1** glycation reactions at 60 °C, extended over several time points (1 h, 2 h, 3 h, 6 h, and 24 h) (n≥3 for all reactions). Under these conditions, [M+144] quickly became the predominant adduct in just 2 h, then decreased precipitously between 2 h and 24 h. During this same time frame, a dramatic increase in APY levels was observed, confirming [M+144] as a precursor to APY. Please note that not all error bars are visible, as many are smaller than the size of the symbol. **(D)** Representative BPCs and MS1 spectra for the glycation reaction of peptide **1** after 24 h incubation at 37 °C (top) and 60 °C (bottom). Together, these results provide conditions at which APY becomes the predominant AGE, suggesting that APY formation likely involves a relatively high energy activation barrier that can be overcome by the addition of heat.

**Figure 4.**
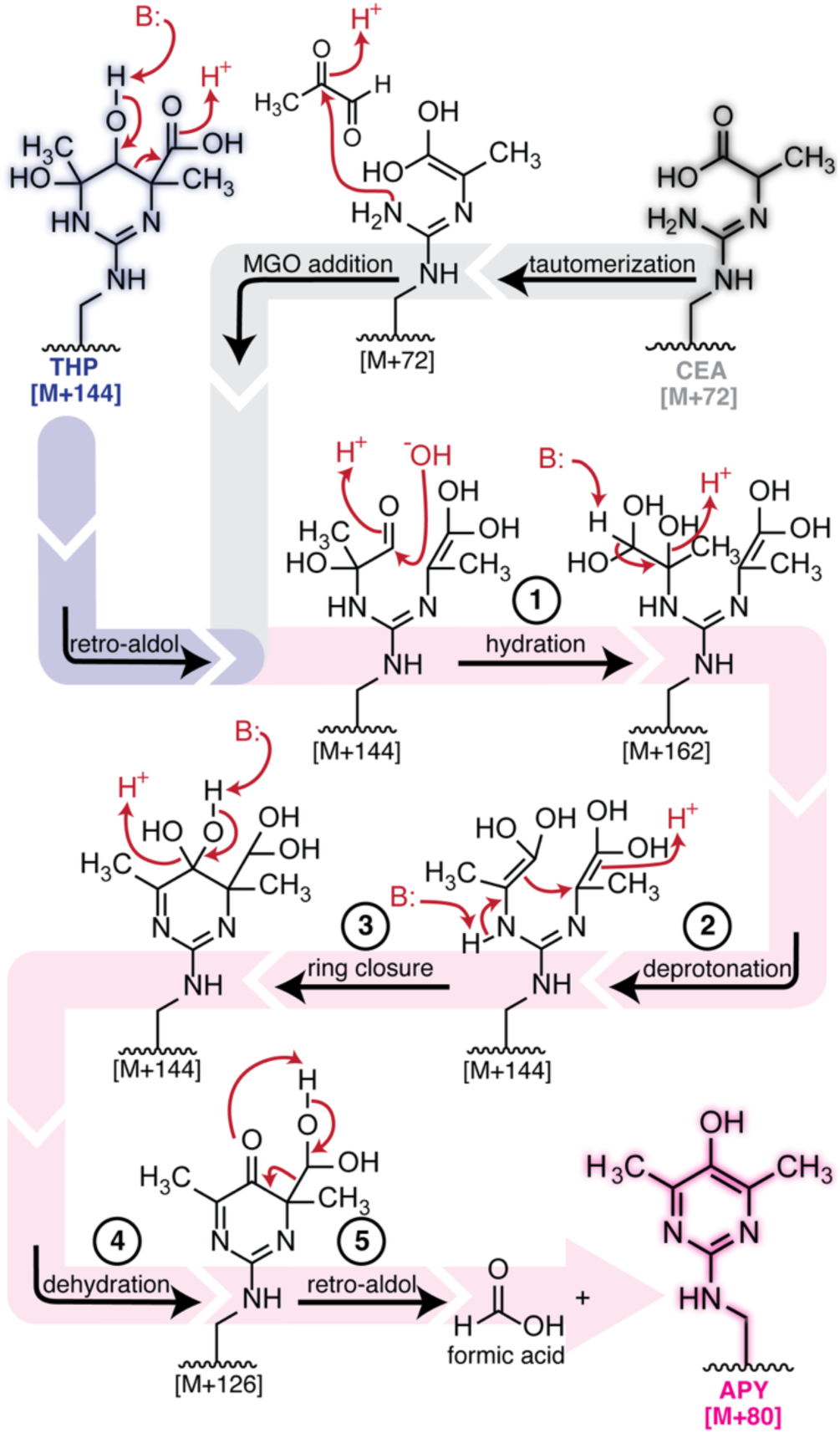
Proposed Mechanism for APY Formation. We propose that THP can rearrange into APY *via* a retro-aldol step (1, *blue path*), followed by hydration (2), then deprotonation (3) and ring closure (4), another dehydration step (5), and finally a retro-aldol step (6) that also releases formic acid as a byproduct (*pink path*). Alternatively, it is likely that other [M+144] species could also rearrange into APY. Here we propose an alternate pathway, beginning from CEA (*gray path*), which involves tautomerization and addition of a second MGO equivalent to reach a common [M+144] intermediate along the APY formation pathway.

### Enhancing APY formation

Despite defining [M+144] as an intermediate along the APY formation pathway, persistent low levels of APY remained a consistent technical challenge. Seeking to increase APY levels, we conducted a temperature scan in which we incubated peptide **1** with 2 mM MGO for 24 hours, using temperatures ranging from 37–70 °C (**Figure 3**). We found that between 37 °C and 45 °C, there was hardly any change in the AGE distribution. Across the entire range of temperatures used, other AGEs, like MGH-1 and CEA, stayed fairly constant, with only slight increases at higher temperatures. However, at 55 °C, we began to observe a notable increase in APY that was concomitant with reduced [M+144] levels. APY levels peaked at 60 °C (53.1 ± 12.1 %), becoming the predominant AGE in just 24 hours (**Figure 3A,B**). Due to high levels of APY observed at 60 °C, we were able to confirm its identity using its characteristic fluorescence wavelengths (ex = 320 nm, em = 385 nm) (**Figure S3**).^2,20,25,27^

To further investigate the relationship between APY and [M+144], we extended the reaction at 60 °C across multiple time points. At 60 °C, the reaction quickly reached 88.1 ± 1.9% total glycation in only 3 h. From this, we conclude that most of the free MGO is quickly consumed, producing [M+144] as the major early AGEs (54.3 ± 1.8% of total glycation) (**Figure 3C**). APY levels began to increase after 2 h of incubation, with a precipitous increase (from 0.0 % to 24.5 ± 5.5 %) between 3 and 6 h. During this same time frame, a concomitant, proportional decrease in [M+144] levels was observed, providing unambiguous assignment of [M+144] as a precursor to APY. Notably, all of the [M+144] species we observed appear to mature into APY, since each one decreased at the same rate as APY levels increased (**Figure S2**). For this reason, we report all [M+144] species totaled together, represented as a combined [M+144].

Additionally, comparison of the AGE distributions at 37 °C and 60 °C revealed APY is a stable AGE. After heating at 60 °C for 24 h, only MGH-1 and APY remained in appreciable quantities, while other AGEs, like CEA and [M+144] species including THP, disappeared, presumably due to rearrangements into MGH-1 and APY (**Figure 3D**). We have also shown that while elevated temperatures increase APY levels at earlier time points, APY formation is still dependent on the sequence identity (**Figure S4**). Thanks to high levels of APY observed, we also conducted stability studies for purified APY on peptide **1** (peptide **1^APY^**) and showed that APY on a short peptide is stable, exhibiting minimal decay for up to a week at physiological conditions (**Figure S5**). Importantly, these findings not only identify conditions at which APY can become a major AGE, but they also suggest that the rearrangement of [M+144] into APY likely involves a high energy activation barrier that can be overcome by the addition of heat.

### A Proposed Mechanism for APY formation

Next, we sought to understand possible mechanisms through which this rearrangement could take place. The most cited APY formation mechanism involves the generation of a reductone from two MGO, which then reacts with Arg to form APY (**Figure S6**).^2^ In this model, preincubation of MGO alone would be expected to generate the reductone, consequently leading to faster APY formation once added to peptide. We found that preincubation, even at strongly acidic pH and elevated temperatures (60 °C), did not promote any further APY formation upon addition to peptide **1** (**Figures S7 & S8**), and we did not observe any evidence of reductone formation by NMR (**Figure S9**). These results are therefore inconsistent with reductone formation as a plausible model for APY formation in physiological conditions. Another proposed mechanism suggests that MGH-3 rearranges to form APY, presumably through oxidative decarboxylation.^24^ However, our previous work shows that peptide **1** preferentially forms the MGH-1 isomer as confirmed by NMR, which we also confirmed in this study by matching the retention time (**Table S1**). As peptide **1** still forms APY, this suggests that MGH-3 is not a prerequisite for APY formation.^4^

Previous proposals have hypothesized that THP rearranges to APY through a pathway involving oxidation, followed by decarboxylation and then dehydration (**Figure S6**).^24,25^ To evaluate the feasibility of an oxidative decarboxylation from [M+144], we first sought to understand whether APY could form under O_2_-free conditions. We scrupulously degassed all reaction components and used a glovebox to assemble and seal glycation reactions inside an N_2_ sparged (O_2_ < 4 ppm) scintillation vial that was further sealed prior to incubation. The resulting APY levels observed were identical in the presence or absence of oxygen (**Figure S10**), suggesting that a formal oxidation step may not be required. However, our studies were performed using commercial sources of MGO, which may contain impurities. To see if any potential impurities were contributing to APY formation, we synthesized MGO using an established Riley oxidation protocol (**Figure S11**).^28^ We found that there was no difference in APY formation between a freshly synthesized stock of MGO and commercial MGO. We also evaluated the role of buffer and salt concentrations, and found negligible differences (**Figure S12**).

As we could not attribute the source of oxidation required in the previously proposed mechanism, we considered other potential mechanisms in which THP—or other [M+144] species—could rearrange to form APY. Beginning from THP, we identified a possible pathway (**Figure 4**) in which initial deprotonation of the hydroxyl group at C5 promotes a retro-aldol ring opening (**Figure 4**, *blue path)*. Next, hydration of the aldehyde followed by deprotonation generates a geminal enediol at C5, followed by subsequent ring closure and another dehydration and retro-aldol step to generate the pyrimidine ring (**Figure 4**, *pink path*). This proposed mechanism generates formic acid as a byproduct. Supporting this proposal, we used an enzyme coupled assay to detect formic acid in reactions of peptide **1** with 2 mM MGO after 24 h (**Figure S13**). Our results indicate that formic acid is a byproduct of APY formation, as more formic acid was detected as APY formed on peptide **1** under our standard glycation conditions. However, we also found that formic acid generation is not necessarily exclusive to APY, since we also detected formic acid when MGO was incubated with another sequence, (Ac-LDDREDA), that exhibited no detectable APY formation.

Our proposed mechanism involves an intermediate that includes a tautomer of CEA on one of the Arg Nω, allowing us to hypothesize that CEA could also be a precursor to APY. To evaluate this idea, we purified CEA and MGH-1 modified peptide **1** (peptides **1^CEA^** and **1^MGH-^**^1^, respectively). At pH 7.3, without any additional MGO added, we observed only modest conversion from **1^MGH-^**^1^ to **1^CEA^** with little degradation to peptide **1**, consistent with our previous results.^4^ Since peptide **1** does not form MGH-3 under our standard glycation reaction conditions, we do not rule out the possibility that MGH-3 can rearrange to form APY. However, upon treatment with additional equivalents of MGO, we found that peptide **1^CEA^** produced APY (**Figure S14**). This suggests that another source of the [M+144] precursor to APY could be due to an additional MGO modification of CEA. This alternate pathway (**Figure 4**, *gray path*) shares a common intermediate with the one that formally begins with THP.

### APY formation requires a general base

To further evaluate the feasibility of our proposed mechanism, we predicted that a general base would be necessary for the initial deprotonation step that promotes ring opening, as well as for subsequent ring closure and dehydration (**Figure 4**). Using a pH scan, we found that APY levels increased between pH 7.3–11 (**Figure 5A,B**). There was a particularly large jump in APY levels between pH 10 (3.3 ± 1.4 %) and pH 11 (20.3 ± 1.1 %), prompting us to conduct a narrower pH scan at 0.2 increments between pH 10 and 11. In this range, we found that APY levels increased steadily, peaking at pH 11 (20.3 ± 1.1 % of total glycation) (**Figure 5C & Figure S15**). There was a sharp transition between pH 10.2 and 10.4, suggesting a strong correlation with the protonation state of the nearby Tyr residue (p*K*_a_ 10.3).^29–31^ As a control, we also synthesized a variant of peptide **1** that replaced Tyr with Phe (peptide **1^Phe^**). Compared to peptide **1**, peptide **1^Phe^** produced substantially less glycation overall and less APY formation (**Figure S16**). These results suggest that, for peptide **1**, Tyr plays an active role in the APY formation mechanism.

**Figure 5.**
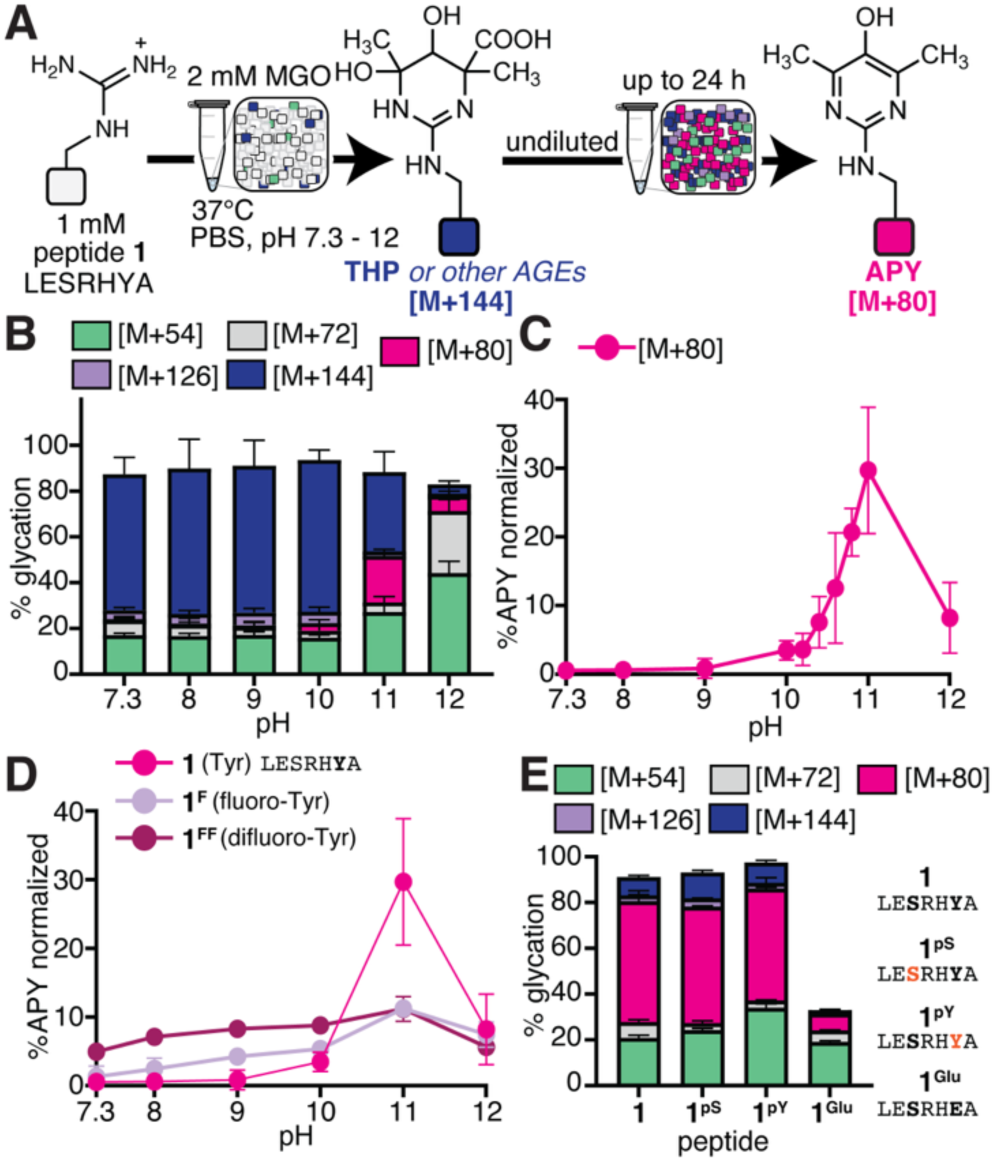
A Nearby General Base Facilitates the Rearrangement of [M+144] into APY. **(A)** General scheme depicting a pH scan used to evaluate the features that promote APY formation. Peptide **1** (1 mM) was treated with 2 mM MGO using pH-adjusted phosphate buffered saline (PBS) in a pH range of 7.3—12. The reaction was incubated for 24 h undiluted at 37 °C. **(B)** Stacked bar graphs showing AGE distributions, as assessed by liquid chromatography-mass spectrometry (LC-MS) analysis. **(C)** Between pH 10 and 11, we observed a marked jump in APY [M+80] (magenta) levels, as shown by proportional APY levels that were normalized to the total amount of glycation observed for MGO-treated peptide **1** across a narrower pH range between pH 10 and 11 at 0.2 increments. **(D)** To further test the hypothesis that Tyr phenoxide plays an active role in APY formation, we conducted glycation reactions on variants of peptide **1**, including peptide **1^F^** (fluoro-Tyr in lieu of Tyr) and peptide **1^FF^** (difluoro-Tyr in lieu of Tyr). Here, proportional APY levels were normalized to the total amount of glycation to account for differences in total glycation. At pH 7.3 and 37 °C, % APY was the greatest for peptide **1^FF^**(p*K*_a_ 7.2), followed by peptide **1^F^** (p*K*_a_ 8.4), and then peptide **1** (p*K*_a_ 10.3), consistent with our hypothesis. **(E)** Stacked bar graphs showing AGE distributions for glycation reactions with peptide **1**, peptide **1^pS^** (pSer in place of Ser, p*K*_a2_ 5.6), peptide **1^pY^** (pTyr in place of Tyr, p*K*_a2_ 5.9), and peptide **1^Glu^** (Glu in place of Tyr) at pH 7.3 and 60 °C (24 h incubation) (n≥3 for all reactions). Despite the introduction of up to two negative charges which has been reported to hamper glycation, absolute APY levels and total glycation levels remained virtually identical for the phosphorylated variants. Together, these results demonstrate the importance of a nearby general base to facilitate APY formation and suggest that nearby phosphorylated residues can also aid glycation.

To test the hypothesis that Tyr phenoxide facilitates APY formation, we synthesized variants of peptide **1** that incorporated noncanonical Tyr derivatives with lowered phenolic p*K*_a_’s. Specifically, 3-fluorotyrosine (peptide **1^F^**) and 3,5-difluorotyrosine (peptide **1^FF^**) were chosen because of their p*K*_a_ values (8.4 and 7.2, respectively) (**Figure 5D**).^32^ These peptides were reacted with 2 mM MGO across a range of pH between 7.3 and 12 (**Figure 5D & Figure S15**), revealing maximal APY levels at pH 11 for all three peptides. However, when normalized to account for differences in total glycation levels, we found that proportional APY levels at pH 7.3 were greatest for peptide **1^FF^** (5.0 ± 0.3%), followed by **1^F^**(1.3 ± 1.5%), and then peptide **1** (0.6 ± 0.4%) (**Figure S15**). These results demonstrate the importance of the nearby Tyr as a facilitator for [M+144] rearrangement into APY, matching our hypothesis that Tyr can act as a general base that facilitates APY formation.

Following the same logic, we next considered if phosphorylated Tyr—or perhaps even Ser or Thr— might also act as a general base promoting APY formation. Phosphorylated Tyr has experimental p*K*_a2_ values ranging between 4.0 to 6.1.^33^ Thus, we synthesized peptide **1^pY^** that contains pTyr in place of Tyr. After glycation with 2 mM MGO at pH 7.3 at 60 °C for 24 h, there were hardly any differences in total glycation and APY levels between peptide **1** (52.8 ± 4.5 % [M+80] at 24 h) and **1^pY^** (48.8 ± 0.6 % [M+80] at 24 h) (**Figure 5E & Figure S17**). We performed a similar set of experiments using a peptide **1** variant with phosphorylated Ser (peptide **1^pS^**), and found that these also produced comparable levels of glycation (50.9 ± 0.3%) after 24 h, despite the addition of up to two negative charges (**Figure 5E & Figure S17**). By contrast, a variant of peptide **1** that replaced Tyr with Glu (peptide **1^Glu^**) exhibited substantially less overall glycation and APY formation than peptide **1** (**Figure 5E & Figure S16**).

To further examine this behavior, we evaluated the glycation of peptides **1**, **1^pY^**, **1^pS^**, and a doubly-phosphorylated version (peptide **1^pSpY^**) at 37 °C, over time (**Figure S18**). Supporting our findings at 60 °C, these studies showed APY levels to be similar across all times for peptides **1**, **1^pY^**, and **1^pS^**. However, MGH-1 levels were noticeably increased—especially at earlier times—for the phosphorylated substrates. Additionally, introduction of up to two negative charges on peptide **1^pY^** and **1^pS^** did not affect overall glycation but led to increased MGH-1 formation. However, at both lower and elevated temperatures, the introduction of four negative charges on peptide **1^pSpY^** decreased overall glycation (**Figures S17 & S18**). Our previous work has unambiguously shown that multiple negative charges surrounding glycation sites greatly impair AGE formation (**Figure 5E**).^4,5,34^ Therefore, this observation suggested to us that nearby phosphates may exert a positive influence on AGE formation, thereby counteracting any attenuation from their negative charges. Accordingly, these results imply that nearby phosphorylation may be helpful for glycation, and based on our proposed mechanism, could be particularly important for APY formation in a cellular context.

### Uncovering a connection between cellular glycation and phosphorylation

Guided by these chemical mechanistic insights, we performed a quantitative proteomics experiment using HEK-293T cells treated with or without MGO. Our mechanistic studies have shown that APY can accumulate at longer times or using higher MGO concentrations. Therefore, we opted to treat cells with 2 mM MGO for 24 h to ensure that we could induce APY formation. We confirmed that these treatment conditions did not significantly decrease cell viability, as more than 95% of cells remained viable after treatment (**Figure S19**). Several reagents, such as dithiothreitol (DTT) and iodoacetamide, used in proteomic experiments were tested for their effects on AGE distribution and APY formation and found to have no interference (**Figure S19**). Additionally, as there are no highly specific antibodies available to enrich APY (**Figure S20**), we chose to label an unenriched cell lysate with isobaric tandem mass tags, allowing us to evaluate the distribution of multiple AGEs using quantitative bottom-up proteomics, including the MGH isomers ([M+54]), APY ([M+80]), THP or other [M+144] AGEs, and an [M+72] adduct that could correspond to MGH-DH, CEA, or CEL. Using this approach, we identified 7210 total proteins and 56490 peptides across all conditions tested, and found 478 unique peptides that were AGE-modified upon treatment with 2 mM MGO. This dataset shows significant similarity to other MGO-treated proteomic datasets, with AGE modifications enriched on proteins involved in processes such as protein folding and spliceosome function, along with several modified histone and heat shock proteins.^9,10,35,36^

Upon meticulous inspection of our dataset, we manually removed any hits from the glycated set that had a Δ mass (theoretical – observed mass) > 20 ppm and/or were assigned with a modified Arg at their C-terminus, as we and others have previously shown that glycated Arg cannot be cleaved by trypsin.^5,37^ After data processing, this decreased the total list of AGE-modified peptides to 204 plausible AGE-modified hits. Without any enrichment steps during sample preparation that could bias the AGE distribution, we found that APY and [M+144] species were the predominant AGEs in the MGO-treated condition (57 unique peptides, 28%, and 53 unique peptides, 26%, respectively), followed by the MGH isomers and CEA/MGH-DH (47 unique peptides, 23% for each) (**Figure 6B**). Although our dynamic modification for [M+72] (mass change 72.021) was set for both Arg and Lys, we only observed modification on Arg (MGH-DH and/or CEA, not CEL), likely due to the static TMT modification that was set for Lys in our processing workflow. Though glycation is largely driven by chemical microenvironments that include 3D protein structure and is therefore unlikely to have a strong consensus sequence,^34^ we were also able to obtain consensus features for each AGE (**Figure S21**).

**Figure 6.**
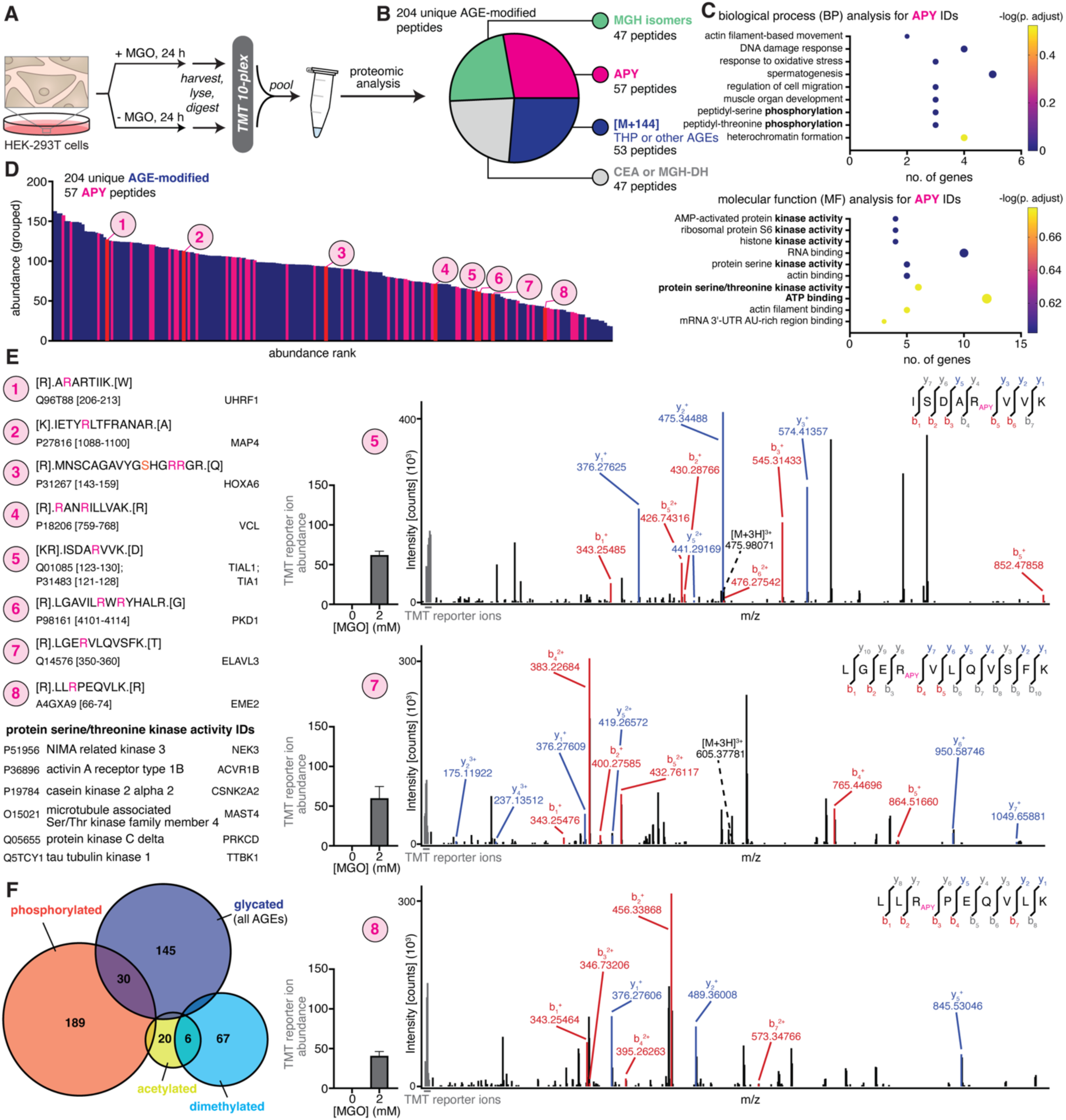
Proteomic Analysis on MGO-treated Cells Reveal a Connection between Glycation and Cellular Phosphorylation Events. Given that our proposed mechanism for APY formation was supported by *in vitro* peptide data, we set out to investigate APY formation in a cellular context, along with other common MGO-modified AGEs. **(A)** General scheme describing our proteomic workflow on HEK-293T cells treated with or without 2 mM MGO for 24 h. Whole cell lysates were trypsin-digested, labeled with TMT isobaric tags, and analyzed in a quantitative bottom-up workflow. The untreated sample was used as a frame of reference to identify any AGE increase in the treated condition. **(B)** Pie chart showing a set of 204 unique peptides with AGE-modified Arg in the MGO-treated condition, consisting of 57 APY-, 53 THP- (or other [M+144] AGEs), 47 MGH-, and 47 CEA/MGH-DH-containing peptides. **(C)** DAVID Gene Ontology (GO) analysis was performed for protein IDs associated with the APY set using a 0.05 false discovery rate (FDR). Values for -log(p-adjust) were calculated using reported Benjamini values from DAVID. Both the biological process (BP) and molecular function (MF) analyses revealed phosphorylation-related terms (bolded text) that are unique to the APY set. **(D)&(E)** The abundance (grouped) for APY-containing peptides (magenta) was ranked and displayed atop the abundance of other AGE-containing peptides (blue). Eight statistically significant increased APY-containing peptides (red) are numbered 1-8. Protein IDs from the protein serine/threonine kinase activity term are also shown in **(E)**. Representative b/y ion spectra and TMT reporter ion abundance quantitation for three of the high confidence APY hits are shown. **(F)** Venn diagram showing overlapping IDs between AGEs or other common Arg modifications (dimethylation and acetylation) and phosphorylation. Compared to dimethylated and acetylated IDs, glycated IDs exhibited substantial overlap (30 IDs) with phosphorylated IDs, suggesting a connection between AGE formation and phosphorylation events.

Next, we performed gene ontology (GO) analysis for the APY-modified set of proteins. In both the biological process (BP) and molecular function (MF) annotations, APY-containing proteins showed enrichment of terms related to phosphorylation, including protein phosphorylation (BP) and multiple kinase activities (MF), despite having a relatively low number of protein IDs (**Figure 6C**). None of the other AGE sets showed an enrichment of protein phosphorylation (BP) or kinase activity (MF) (**Figure S22—S25**), and the APY-modified set was the only set to exhibit enrichment for protein serine/threonine kinase activity, implying a connection between APY formation and phosphorylation in a cellular context.

Building on this observation, we further investigated the set of 57 modified APY-containing peptides, which were observed across the entire range of AGEs when ranked by abundance (**Figure 6D**). We focused in particular on the eight highest confidence APY-modified peptides that exhibited statistically significant increases in abundance upon MGO treatment across all three biological replicates (**Figure 6D,E** and **Figures S26—S30**). Data imputation and statistical analysis were performed according to published protocols.^38^ Among these peptides, one was also identified with a phosphorylated residue nearby the APY-modified site, corresponding to HOXA6 (hit 3) (R156, R157 with nearby pS153).

By cross-referencing the protein sequences, reported phosphorylated sites, and the influence of phosphorylation on protein functions, we were able to draw several correlations between reported studies and our dataset. Of the remaining seven high confidence APY hits, six were previously reported to be phosphorylated, and some are known to be regulated by phosphorylation.^39^ For example, phosphorylation of MAP4 (hit 2) has been reported to affect microtubule properties and cell cycle progression, while TIAL1 (hit 5) has also been found to be phosphorylated following DNA damage, and phosphorylation of PKD1 (hit 6) triggers its membrane dissociation and subsequent entry into the nucleus.^40–42^ Additionally, phosphorylation of UHRF1 maintains protein stability, subcellular localization, and epigenetic regulation (hit 1).^43^ Careful inspection of our data revealed that UHRF1 was APY-modified at R207, which is close to T210, a known phosphorylated site previously reported to be essential in maintaining UHRF1 stability, though phosphorylated T210 was not observed in our dataset.^44^

Similarly, S358, close in sequence to APY-modified R353, on ELAVL3 (hit 7) has also been shown to be phosphorylated.^39^ We also found two APY modifications on the C-terminal domain of PKD-1 (R4107, R4109), which contains 4 Tyr residues (Y4110, Y4118, Y4127, Y4237) and 2 putative Ser sites (S4169, S4252) that are susceptible to phosphorylation. Multiple *in vitro* studies have revealed Y4237 and S4252 to be targets of phosphorylation by pp60c-src and PKA, respectively, and shown endogenous PKD-1 to be phosphorylated at tyrosine.^45–47^ Together, our data show several APY modifications on Arg residues close to known phosphorylation sites, consistent with the potential for crosstalk between cellular APY and phosphorylation events.

However, as APY (mass change 80.026) is quite close in mass to phosphorylation (mass change 79.966), we also queried our dataset for phosphorylation and found that the list of proteins generated had some overlap with the set of APY-modified targets, but the overall list of IDs was quite different, confirming that the relatively similar mass changes did not produce artifactual hits. We also found that the phosphorylated set (220 proteins) showed substantial overlap (30 proteins) with the AGE-modified set of proteins (180 proteins). To evaluate the possibility that crosstalk between nearby phosphorylation could promote AGE formation (especially for APY and MGHs), we performed a parallel analysis for other common Arg PTMs, including dimethylation and acetylation (**Figure 6F** and **Figures S31 & S32**). We found 99 unique peptides with dimethylated Arg and 30 unique peptides with acetylated Arg. Importantly, the phosphorylated set of proteins did not substantially overlap with those that were observed to be dimethylated (9 IDs) or acetylated (2 IDs) (**Figure 6F**).

### Confirming that phosphorylation can potentiate glycation

Overall, our proteomic dataset supported our *in vitro* experiments, suggesting that APY formation might be promoted by phosphorylated residues that act as general bases to facilitate rearrangement into APY **(Figure 4)**. To validate our findings, we synthesized three peptides matching APY-modified sequences in our dataset. These include a sequence from MAP4 (hit 2; peptide **V1**: Ac-ETYRLTF), and one from ELAVL3 (hit 7, peptide **V2**: Ac-LGERVLQ). We also synthesized a peptide from HNRNPA3, which was found in higher abundance than either hit sequence, though it did not meet our threshold for being considered “high confidence” (peptide **V3**: Ac-YNLRDYF). Upon treatment with 2 mM MGO at 60 °C for 24 h, we confirmed that all three peptides formed APY (**Figure 7A,B**). Expectedly, peptide **V2**, without a nearby Tyr, exhibited the least APY formation out of the three peptides, though we note that it produced more APY than control sequences such as peptide **1^Glu^**and peptide **1^Phe^**. Matching our expectations, we found that the peptide with two Tyr residues (peptide **V3**) exhibited the most APY formation. We also found that the overall extent of glycation and the amount of APY formed for peptide **V1** and **V3** were quite similar to that observed for peptide **1**. However, as our treatment condition at 60 °C for 24 h does not reflect the initial rate of APY formation, we also monitored the AGE distributions at 37 °C at 24 and 48 h (**Figure 7C & Figure S33)**. Under these conditions, peptide **V1** exhibited the most APY formation. This behavior was visible at earlier time points, and was particularly apparent after 48 h of treatment. Importantly, these data validate that peptides matching the APY-modified sequences identified in our proteomic analysis are indeed capable of forming APY *in vitro*.

**Figure 7.**
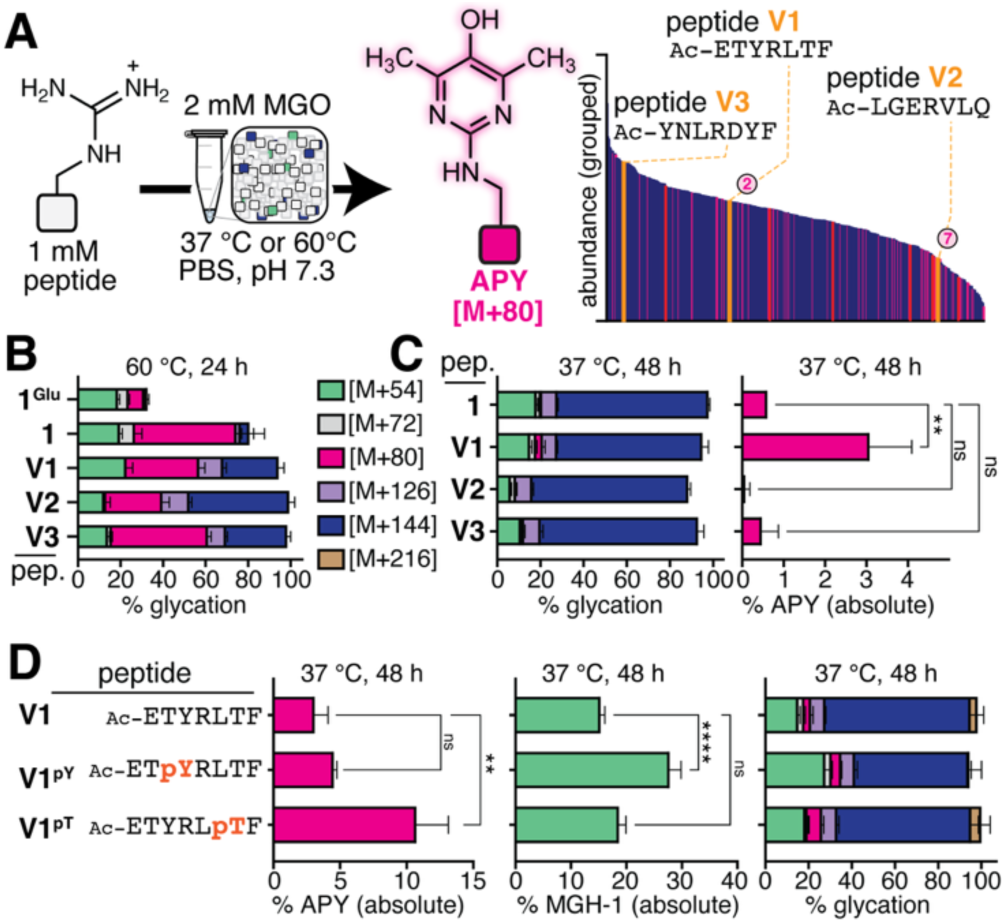
Confirming that phosphorylation can potentiate APY formation *in vitro*. **(A)** To validate our proteomics analysis, we synthesized peptides matching APY-modified hit sequences in our dataset: Ac-ETYRLTF (hit 2; peptide **V1**), Ac-LGERVLQ (hit 7; peptide **V2**), and Ac-YNLRDYF (peptide **V3**), a high-abundance sequence. These peptides were evaluated after treatment with MGO **(B)** Distributions of AGEs at 60 °C after 24 h of MGO treatment for peptides **1**, **V1**, **V2**, **V3,** and **1^Glu^** (n = 3). All three peptides identified from proteomic analysis were confirmed to form APY. **(C)** Distributions of AGEs at 37 °C after 48 h of MGO treatment (*left*) for peptides **1**, **V1**, **V2**, and **V3**. Absolute APY levels (%) observed under the same conditions (*right*) (n = 3) shows that peptide **V1** exhibits the most APY formation. Ordinary one-way ANOVA was used to determine if each variant yielded statistically significant differences in % absolute APY compared to peptide **1**. p<0.01(**). **(D)** Absolute APY (*left*) or MGH-1 (*middle*) levels, and full distributions of AGEs (*right*) after 48 h of MGO treatment at 37 °C for peptides **V1**, **V1^pY^**, and **V1^pT^** (n = 3). All three peptides exhibited similar levels of overall glycation. Peptide **V1^pT^** was the fastest to form APY, followed by peptide **V1^pY^**, and then peptide **V1**. Ordinary one-way ANOVA was used to determine if each variant yielded statistically significant differences in APY or MGH-1 levels compared to peptide **V1** (n = 3). p<0.01(**), p<0.0001(****). Absolute APY levels in peptide **V1^pT^** were three-fold higher than those in peptide **V1**, and both phosphorylated variants exhibited an increase in MGH-1 formation, indicating that phosphorylation has a direct influence on glycation outcomes.

As peptide **1** was identified from a prior study focused on MGH-1, we found that introduction of nearby phosphates appeared to have an impact on MGH-1, rather than APY, formation. To determine if this was also the case for hit peptides, we synthesized two variants of peptide **V1** that incorporated either pTyr or pThr (Ac-ETpYRLTF, peptide **V1^pY^**, and ETYRLpTF, peptide **V1^pT^**) (**Figure 7D & Figure S33**). At 60 °C for 24 h, all three peptides exhibited similar levels of overall glycation and APY formation, despite the introduction of extra negative charges. Compared to peptide **V1^pY^**and non-phosphorylated peptide **V1**, peptide **V1^pT^** showed a significant increase in MGH-1 formation. However, at 37 °C, peptide **V1^pT^**was the fastest to form APY, followed by peptide **V1^pY^**, and then peptide **V1** (**Figure 7D**). Absolute APY levels in peptide **V1^pT^**were three-fold higher than those in peptide **V1**. Interestingly, similar to the trends observed for peptide **1** and its phosphorylated variants, peptide **V1^pT^** and **V1^pY^** also exhibited an increase in MGH-1 formation, with absolute MGH-1 levels in peptide **V1^pY^** observed to be approximately two-fold higher than those in peptide **V1** (**Figure 7D, Figure S33**), indicating that phosphorylation can also bias the formation of certain AGEs. Taken together, these data support a model in which crosstalk with protein phosphorylation could have an important, but previously underappreciated role in regulating cellular glycation events.

## Discussion

Herein we have used a peptide model system to determine a chemical mechanism for APY formation. Although appreciated for its unique fluorescence properties,^2,24,48^ APY has been difficult to study as it was not previously known how to produce it in experimentally convenient quantities from the reaction of Arg and MGO.^17,18,20^ Our results indicate that APY formation depends surprisingly little on initial MGO concentrations, and instead greatly depends on other factors, including sequence identity, reaction pH, and temperature, each of which can help to overcome a relatively high energy rate determining step. Through these studies, we discover that APY is quite stable and identify *in vitro* conditions under which APY becomes the predominant AGE, which to our knowledge, has not been previously reported. These findings alone provide an important technical advance that will greatly facilitate future studies on APY.

Additionally, our proposed mechanism relies on fundamental concepts of organic chemistry and provides concrete experimental evidence that reconciles several previous, conflicting mechanisms proposed for APY. While others have suggested that THP is a precursor to APY, our newly proposed mechanism circumvents the need for an unexplained, mysterious oxidation step by recognizing that THP possesses a β-hydroxycarbonyl, a common feature in retro-aldol substrates. Indeed, the retro-aldol reaction is one of the most important organic reactions in sugar fragmentations, including those in physiological systems.^49–51^ Several known retro-aldol enzymes use Tyr as a general base,^52–54^ which aligns with our results showing noncanonical Tyr variants with decreased phenolic p*K*_a_ to promote APY formation. However, we also identify a potential alternative pathway involving highly plausible [M+144] intermediates other than THP. While we cannot definitively confirm all steps in our proposed mechanism, it is fully consistent with our experimental results ruling out other previously proposed alternatives. Additionally, we note that most steps are precedented in biological systems, including the retro-aldol reaction^49–51^ and tautomerization of hydrated aldehyde consistent with glyoxalase activity.^55–58^ Our mechanism also connects APY formation to the emancipation of formate, a major reactive species in one-carbon metabolism that is involved in numerous biological processes.^59–61^ While there has been some prior work connecting the metabolic processing of MGO to formate generation,^62–66^ our results suggest that cellular glycation events themselves, including but not limited to APY formation, may also contribute to the cellular formate pool.

Encouragingly, our dataset shows that APY is likely to be a major AGE in cells. We suspect the prevalence of APY has seemingly been underestimated in previous studies, either due to the use of enrichment protocols that are poorly suited to capture APY,^67,68^ or due to a focus on AGEs that were previously established to be abundant in prior work.^69,70^ Our mechanistic studies therefore not only provide much needed insights into APY formation under physiologically relevant conditions, but also reveal critical aspects of the glycation landscape. In future studies, we plan to leverage this information to generate novel chemical tools that can be used to track and/or control APY formation in living cells, including the development of tools that harness APY’s unique autofluorescent properties for bioconjugation and to assist in the study of APY biology.

This study also reveals the potential for significant crosstalk between glycation and phosphorylation, showing the presence of nearby phosphorylated side chains to enhance, and potentially bias, particular AGEs to form. Our experiments examine the ability of phosphorylated residues (pSer, pTyr, & pThr) to promote formation of certain AGEs, particularly that of MGH-1 and APY, on nearby Arg despite introducing additional negative charges previously shown to hamper glycation.^4,5^ These peptide data are also supplemented with quantitative proteomics analysis on MGO-treated cells, revealing a similar connection between phosphorylation and glycation, that was notably missing with other Arg PTMs like acetylation and dimethylation. While inorganic phosphate has long been known to enhance glycation,^71^ and MGO has been shown to stimulate phosphorylation,^72–75^ we have not been able to find any reports suggesting that nearby phosphorylated sites influence glycation. Therefore, our study is the first to show that glycation may be coincident with phosphorylation on the same substrates, and that the presence of these nearby phosphates may bias the resulting AGE distribution. Our future work will seek to fully elucidate both the chemical and biological significance of this phenomenon.

## ASSOCIATED CONTENT

### Supporting Information

All AGE peptides from proteomic analysis of treated cells in Excel file

### Author Contributions

V.T.T.P. designed the project, performed all peptide experiments and analysis, analyzed proteomics datasets, and prepared the figures and manuscript. S.D. acquired and assisted in the proteomics data analysis. A.C.S assisted in cell culturing and viability testing and synthesized MGO using protocols developed by S.M.H. R.A.S designed the project, directed the experimental design and analysis, and prepared the figures and manuscript. All authors have given approval to the final version of the manuscript.

### Funding Sources

This work was supported by National Institutes of Health grant R01GM132422 to R.A.S, as well as by a gift to the Scheck Lab from J. Kanagy, and a gift to the Scheck Lab from J. Fickel.

## ACKNOWLEDGMENT

We would like to express our gratitude to Dr. Krishna Kumar and Dr. Luke Davis for project guidance and to the Zhang Lab (Tufts University Department of Chemistry) for assistance with experiments conducted inside the glovebox.

## ABBREVIATIONS

AGE: advanced glycation end-product
APY: argpyrimidine
MGO: methylglyoxal
MGH-1: -2, & -3, hydroimidazolone isomers
MGH-DH: dihydroxyimidazolidine
CEA: carboxyethylarginine
THP: tetrahydropyrimidine
LC-MS: liquid chromatography-mass spectrometry
HPLC: high-performance liquid chromatography
TMT: tandem mass tag
GO analysis: gene ontology analysis.

