## Supporting Information for "A Chemical Mechanistic Path Leads the Way to Cellular Argpyrimidine"

**Materials and Methods** 2

**Supplementary Figures and Tables**

**Figure S1.** Peptide Glycation for Previously-Identified APY-modified Protein Sequences 7

**Figure S2.** All [M+144] Species Mature into APY over Time 8

**Figure S3.** Detection of APY Fluorescence by Analytical HPLC 9

**Figure S4.** APY Formation can be Affected by the Surrounding Sequence 10

**Figure S5.** APY on a Short Peptide is Stable under Physiological Conditions 11

**Figure S6.** Previously Proposed APY Formation Mechanisms 12

**Figure S7.** MGO Preincubation at Different pH does not generate APY 13

**Figure S8.** MGO Preincubation at Elevated Temperatures does not produce APY 14

**Figure S9.** NMR Spectra do not show Reductone Formation by MGO Preincubation 15

**Figure S10.** Oxygen is Not Required to Form APY 16

**Figure S11.** Synthetic and Commercial MGO Sources both generate APY 17

**Figure S12.** Phosphate Buffer and Salt Concentrations do not impact APY formation 18

**Figure S13.** Detection of Formic Acid during Glycation Reactions 19

**Figure S14.** CEA is also a Precursor for APY 20

**Figure S15.** MGO Treatments of Fluorinated Variants of Peptide **1** 21

**Figure S16.** Replacing the Tyr with Glu and Phe on Peptide 1 reduced Glycation and APY Formation 22

**Figure S17.** Phosphorylated Amino Acids bias Certain AGE Formation at Elevated Temperatures 23

**Figure S18.** Phosphorylated Amino Acids bias Certain AGE Formation at Physiological Temperature 24

**Figure S19.** Cell Viability and Controls showing that Proteomic Reagents do not impact the AGE Distribution 25

**Figure S20.** α-MGH-1 and α-APY Antibody Screening for On-bead Peptide Glycation 26

**Figure S21**. Frequency Logos for All AGE-modified Peptides Observed in Proteomic Analysis 27

**Figure S22—S25.** DAVID GO Analysis Annotation Charts for AGE Modifications 28

**Figure S26—S30.** MS/MS Analysis for Additional High Confidence APY Hit Peptides 32

**Figure S31—S32.** DAVID GO Analysis Annotation Charts for Additional Common Arg Modifications 37

**Figure S33.** Proteomic Analysis Validation with in vitro Peptide Data 29

**References** 40

### Materials and Methods

**General Materials.** All chemical reagents and solvents were of analytical grade, obtained from commercial suppliers, and used without further purification, unless otherwise noted. Methylglyoxal (MGO) (40% w/v in water) was purchased from MilliporeSigma (M0252). N-Fmoc-3-Fluoro-L-tyrosine (1270290) was purchased from Acrotein Chem Bio Inc.. N-Fmoc-3,5-Difluoro-L-tyrosine (AS01393)was purchased from Acrotein Chem Bio Inc.. Fmoc-Tyr(PO(OBzl)OH)-OH (FY2580) was purchased from Advanced ChemTech Inc.. Fmoc-Ser(HPO3Bzl)-OH (101604) was purchased from ChemPep Inc.. Aminoguanidine hydrochloride (396494), NAD (10127981001) and NADH (10128023001) standards were purchased from Sigma Aldrich. Formic acid assay kit was purchased from Neogen Corporation (K-FORM). HEK-293T cells were purchased from ATCC. Water used in glycation experiments was distilled and deionized using an Arium Pro purification system (Sartorius). All statistical analysis was conducted using Prism GraphPad.

**Peptide Synthesis.** Peptides were synthesized using standard Fmoc-based solid phase peptide synthesis on Trityl chloride resin (200 mesh, 1.6 mmol/g loading, Novabiochem®) or Fmoc-Ala-Wang resin (100-200 mesh, 0.64 mmol/g loading, Creosalus Inc.), typically on a 100 μmol scale in a 3 mL polypropylene fritted syringe. For Trityl chloride resin only, C-terminal amino acids (5 equiv., 500 μmol) were coupled overnight in a solution of 10 equiv. (1 mmol) N,N-diisopropylethylamine (DIEA) in 3 mL dimethylformamide (DMF). The next day, the resin was washed and capped with a solution (3 mL) of DMF:MeOH:DIEA (17:2:1 by volume) for 1 hour before further deprotection and coupling steps. C-terminal coupling and resin capping was not required when using Fmoc-Ala-Wang resin. Fmoc deprotection was accomplished using 20% piperidine in DMF (3 mL, 3 x 5 min), followed by washing steps with DMF (3 mL, 4 x 1 min). Amino acid couplings were completed by incubation of amino acid (5 equiv. relative to resin loading) with O-(benzotriazole-1-yl)-N,N,N',N'-tetramethyluronium hexafluorophosphate (HBTU, 5 equiv.) and DIEA (10 equiv.) in 1-2 mL of DMF for 45-90 min. To prepare for pTyr and pSer couplings, deprotection steps right before and after each coupling step were extended to 3 x 10 min, and the couplings were extended to 2-3 hours. All peptides were N-terminally acetylated following deprotection of the last amino acid coupling via incubation with acetic anhydride (4 equiv.) and DIEA (3 equiv.) in 1-2 mL DMF for 2 hours. Following the final acetylation step, peptides were washed with DMF (3 mL, 4 x 1 min), followed by dichloromethane (DCM) (3 mL, 2 x 1 min, then 2 x 15 min), and stored under vacuum desiccation until ready for side-chain deprotection and cleavage. The side chain protecting groups used were as follows: Glu (tBu), His (Trt), Lys (Boc), pSer (Bzl), pTyr (Bzl), Trp (Boc), and Tyr (tBu). Side-chain deprotection and peptide cleavage was completed by incubation with trifluoroacetic acid (TFA), triisopropylsilane (TIPS), and water (95:2.5:2.5 by volume) in 4 mL of acid solution per resin for 2 hours. The resulting crude peptide was dried under constant air flow and then dissolved in 1-3 mL of a water/acetonitrile mixture based on solubility, prior to purification.

**Peptide Purification.** All peptides were purified using a semi-preparative Agilent 1260 Infinity LC system equipped with an Agilent ZORBAX SB-C18 column (9.4 × 250 mm, 5 μm particle size). The mobile phase contained water (A) and acetonitrile (B) with 0.1% TFA. Crude peptide solutions were eluted with a gradient of 5% B to 40% B over 20 min and a flow rate of 3.0 mL/min. Crude peptide solutions of pTyr- and pSer-containing peptides were eluted with a gradient of 5% B to 35% B over 23 min and a flow rate of 2.0 mL/min. Absorbance at 215 and 280 nm was used to observe desired peptide peaks, which were then eluted and collected using an automated fraction collector. APY-specific absorbance could be detected using its characteristic absorbance at 320 nm.[1–3] Collected fractions were assessed for purity using matrix-assisted laser desorption/ionization time-of-flight (MALDI-TOF) mass spectrometry (Bruker) and/or an Agilent 6530 quadrupole time-of-fight (Q-TOF) mass spectrometer (Agilent). Pure fractions were combined and lyophilized. Peptide stock solutions were prepared at 20 mM concentrations in DMF. Peptide stock solutions of peptides containing multiple phosphorylated amino acid on the same sequence were prepared at 10 mM concentrations in 10 mM phosphate buffer (pH 10.0).

**MALDI Mass Spectrometry.** 0.5 μL of each peptide collected fraction were co-crystallized onto a ground steel plate with 0.9 μL of saturated solutions of α-cyano-4hydroxycinnamic acid in 50% acetonitrile, 50% water with 0.1% trifluoroacetic acid.

**Liquid Chromatography-Mass Spectrometry Analysis and Quantification.** Reversed-phase liquid chromatography and mass spectrometry (LC-MS) was performed using an Agilent 1260 LC system coupled to an Agilent 6530 Accurate Mass Q-TOF. The mobile phase contained water (A) and acetonitrile (B) with 0.1% formic acid. Peptide glycation reactions were injected onto an AdvanceBio Peptide 2.7 μm column (2.1 × 150 mm, Agilent) with the following method with a flow rate of 0.4 mL/min: isocratic at 5% B between 0.00-1.75 min, gradient change from 5% B to 40% B between 1.75-16.00 min, gradient change from 40% B to 100% B between 16.00-20.00 min, isocratic column washing at 100% B between 20.00-23.00 min, and re-equilibration at 5% B between 23.01-30.00 min. Peptide data were quantified using peak volumes determined by Agilent MassHunter Qualitative Analysis and the MassHunter Molecular Feature Extractor based on cumulative MS ion counts for 'volumes' observed for any and all charge states associated with a particular ion.

Percent glycation was quantified according to the formula:

$$\%\ glycation = \frac{volume\ of\ AGE\ modified\ peptide}{total\ volume\ of\ both\ modified\ and\ unmodified\ peptide}$$

**Table 1.** AGE adducts for peptide 1, reported with exact mass (Da), Δ Mass (Da), and retention time (RT, min)

| | **Exact Mass (Da)** | **Δ Mass (Da)** | **RT (min)** |
|---|---|---|---|
| **unmodified peptide 1** | 916.4368 | 0 | 9.33 |
| **1**$^{\mathbf{MGH\text{-}1}}$ | 970.4482 | 54.0114 | 10.0 |
| **1**$^{\mathbf{MGH\text{-}DH}}$ | 988.4621 | 72.0253 | 9.43 |
| **1**$^{\mathbf{CEA}}$ | 988.4591 | 72.0223 | 10.10 |
| **1**$^{\mathbf{APY}}$ | 996.4646 | 80.0278 | 11.04 |
| **1**$^{\mathbf{[M+144]}}$ | 1060.4814 | 144.0446 | 10.03 |
| **1**$^{\mathbf{[M+144]}}$ | 1060.4818 | 144.0450 | 10.35 |
| **1**$^{\mathbf{[M+144]}}$ | 1060.4822 | 144.0454 | 9.55 |
| **1**$^{\mathbf{[M+144]}}$ | 1060.483 | 144.0462 | 9.90 |

**Peptide Glycation Protocol.** Peptide glycation reactions were carried out in Eppendorf tubes with a final volume of 50 μL. 10 mM MGO stocks were prepared by adding 15.38 μL commercial MGO solution (40% w/v) into 10 mL of ultrapure water and stored at 4°C for up to one week. 100 mM PBS stocks (pH 7.3) were prepared using BupH Phosphate Buffer Saline packs according to instructions provided by Thermo Scientific (28372). Typically, a peptide in vitro glycation reaction contained 27.5 μL of ultrapure water, 10 μL of 10 mM MGO stock in water (2 mM final concentration), 10 μL of 100 mM PBS stock in water (20 mM final concentration), and 2.5 μL of 20 mM peptide stock solution in DMF (1 mM final concentration). Tubes were capped, briefly spun using a benchtop microcentrifuge, and incubated in a 37°C water bath for up to 24 h or up to 6 weeks depending on the experiment. After incubation, reactions were diluted 100X with ultrapure water and 500 mM Tris Buffer pH 7.4 (5 mM final concentration) to quench the reaction, and then subjected to LC-MS analysis.

**Peptide Glycation at Elevated Temperatures.** Peptide glycation reactions were carried out according to the general glycation protocol for final concentrations. During screening, reactions were assembled in PCR tubes and incubated in a thermocycler (Biorad) using a temperature gradient so that each temperature is set for each row (37 °C, 45 °C, 55 °C, 60 °C, 65 °C, and 70 °C). Subsequently, glycation reactions were performed at 37 °C and 60°C with incubation in water baths.

**Peptide Glycation at Different pH.** Peptide glycation reactions were carried out according to the general glycation protocol for final concentrations. 100 mM PBS stocks were prepared using BupH Phosphate Buffer Saline packs according to instructions provided by Thermo Scientific (28372). The pH of each PBS stock was then adjusted with HCl (100 mM) or NaOH (100 mM) to the desired pH (7.3, 8.0, 9.0, 10.0, 10.2, 10.4, 10.6, 10.8, 11.0, and 12.00), monitoring with a benchtop pH meter (VWR).

**MGO Preincubation Protocol for Peptide Glycation.** Prior to preparation of 10 mM MGO stocks, the ultrapure water pH was adjusted to the desired pH (2.0, 7.3, or 11.0), using strong acid calculations to determine the exact amount of 100 mM hydrochloric acid (HCl) or 100 mM sodium hydroxide (NaOH) required. Using the pH adjusted water, 10 mM MGO stocks were prepared as described above. In preincubation studies, each MGO stock was then preincubated at 37 °C or 60 °C for up to 72 hours, and peptide glycation reactions were assembled using these stocks. To ensure that the pH of the glycation reaction was maintained at 7.3, 10 μL of ultrapure water containing the same volume and concentration of HCl (if preincubated MGO stock was adjusted by addition of NaOH) or NaOH (if preincubated MGO stock was adjusted by addition of HCl) was added to adjust the pH of the 10 mM MGO stock solution back to 7.3 prior to assembling the glycation reaction. The final glycation reaction (50 µL final volume) was prepared by adding 10 µL of 20 mM MGO in ultrapure water at pH 7.3 (2 mM final concentration) to 17.5 µL of ultrapure water, 10 µL of pH-adjusted ultrapure water using either HCl or NaOH as described above, 10 µL of 100 mM PBS stock in water (pH 7.3, 20 mM final concentration), and 2.5 µL of 20 mM peptide stock solution in DMF (1 mM final concentration, 5% DMF).

**Preparation of APY-Modified Peptide 1.** APY-modified peptide 1 (peptide $1^{APY}$) was prepared by incubation of 200 µL of 20 mM peptide 1 stock in DMF with 400 µL of 10 mM MGO stock in ultrapure water and 400 µL of 100 mM PBS stock in ultrapure water (pH 7.3) at 60 °C for 24 hours. Under this condition, APY was the predominant adduct (roughly 50% yield by LC-MS) and was purified by semi-preparative HPLC system as described above using a gradient of 5% to 50% acetonitrile (B) in water (A) over 14.25 min at 3.0 mL/min. Collected fractions were characterized by MALDI-TOF, pooled, lyophilized, and assessed for purity by LC-MS prior to further experiments. The identity of peptide $1^{APY}$ was confirmed by its characteristic absorbance at 320 nm and fluorescence excitation and emission at 320 nm and 385 nm, respectively.

**Preparation of MGH-1-Modified Peptide 1.** MGH-1-modified peptide 1 (peptide $1^{MGH\text{-}1}$) was prepared according to established protocol by McEwen et al. by incubation of 200 µL of 20 mM peptide 1 stock in DMF with 400 µL of 10 mM MGO stock in ultrapure water and 400 µL of 100 mM PBS stock in ultrapure water (pH 12) at 37 °C for 3 h.[4] Under this condition, MGH-1 was the predominant adduct (30-50% yield by LC-MS) and was purified by semi-preparative HPLC system as described above using a gradient of 10% to 30% acetonitrile (B) in water (A) over 20 min at 4.0 mL/min. Collected fractions were characterized by MALDI-TOF, pooled, lyophilized, and assessed for purity by LC-MS prior to further experiments. The identity of peptide $1^{MGH\text{-}1}$ was confirmed by retention time on LC-MS.

**Preparation of CEA-Modified Peptide 1.** CEA-modified peptide 1 (peptide $1^{CEA}$) was prepared according to established protocol by McEwen et al. with minor changes.[4] Briefly, 100 µL of 20 mM peptide 1 stock in DMF was incubated with 200 µL of 100 mM PBS stock in ultrapure water (pH 7.3), 500 µL of 100 mM PBS stock in ultrapure water (pH 12), and 700 µL ultrapure water at 37 °C for up to 1 week. Under this condition, CEA was the predominant adduct and was purified by semi-preparative HPLC system as described above using a gradient of 20% to 35% acetonitrile (B) in water (A) over 20 min at 3.5 mL/min. Collected fractions were characterized by MALDI-TOF, pooled, lyophilized, and assessed for purity by LC-MS prior to further experiments. The identity of peptide $1^{CEA}$ was confirmed by retention time on LC-MS.

**General Protocol for Dilution Experiments.** For dilutions experiments described in Main Text Fig. 2, peptides (1 mM) were incubated with MGO (2 mM) as described above. At time points of interest (typically 3 h and/or 24 h), an aliquot of the reaction mixture (10 µL) was diluted into 990 µL 20 mM PBS (pH 7.3, 100X dilution), briefly spun using a benchtop microcentrifuge, and further incubated in a 37 °C water bath for up to 48 additional hours. The resulting diluted reaction mixture was subjected to LC-MS analysis without any further dilution or purification.

**Preparation of Oxygen-free Glycation Reactions.** To perform glycation reactions in inert atmosphere, each reaction component (water, 10 mM MGO stock, 100 mM PBS stock, 20 mM peptide stock) was carefully degassed using $N_2$ for 30 minutes and sealed inside a 20 mL glass screw-neck scintillation vial with vinyl electrical tape prior to transferring inside a VAC-ATM Genesis glovebox filled with $N_2$ gas ($O_2$<4ppm). Glycation reactions were then assembled in Eppendorf tubes inside the glovebox, according to our standard peptide glycation protocols as described above. The tape-sealed tubes were then sealed inside glass scintillation vials, which were sealed with high adhesion, moisture tight vinyl tape to limit air exposure when transferring reactions out of the glovebox and during incubation. Reactions performed at 37 °C reactions were then incubated in a VWR incubator, while those at 60 °C were incubated in a water bath for 24 h. Reactions were prepared for LC-MS analysis by diluting 100X in degassed water inside the glovebox, and sample vials were sealed with tape until ready for injection.

**Methylglyoxal (MGO) Synthesis Protocol.** MGO was synthesized as described by Riley Oxidation.[5] To start, equal parts (10 mmol) acetone and selenium dioxide powder were added to water to a total volume of 5 mL in a round bottom flask with a magnetic stir bar. The reaction was heated in an oil bath at 100 °C under reflux for 4 hours. After reflux, distillation was performed by attaching the flask to a short path distillation head, and the mixture was distilled into one fraction at an internal temperature of 100 °C. The pale-yellow distillate was lyophilized overnight, revealing a viscous yellow MGO product (134 mg, 19%). Synthetic MGO was characterized by NMR, and a synthetic MGO stock was prepared by adding ultrapure water and checked for concentration using an aminoguanidine assay.[6,7] After a 5-hour incubation of aminoguanidine and MGO at 37 °C, absorbance was measured at 320 nm to generate a standard curve, the slope of which was compared to that of a calibration curve generated by serial dilutions of commercial MGO stock to equalize MGO concentrations between synthetic and commercial stocks.

**NMR Acquisition.** $^{1}$H and $^{13}$C spectra were acquired with a Bruker Advance III (500 MHz, 125 MHz) spectrometer. The following convention was used in data reporting: chemical shift, multiplicity (s = singlet, d = doublet, t = triplet, q = quartet, m = multiplet), coupling constants (Hz), and integration, if applicable. $^{1}$H $D_2O$ δ 5.25 (s, 1H), 4.79 (s, 1H), 2.28 (s, 3H), 1.35 (s, 3H). $^{13}$C (126 MHz, $D_2O$) δ 209.13, 95.13, 91.89, 89.72, 24.53, 21.40.

**Formic Acid Assay.** Formic acid was detected using a formic acid assay kit purchased from Megazyme (K-FORM). This assay utilized the enzyme formate dehydrogenase (FDH) to detect formic acid in samples by converting $NAD^+$ to NADH. The absorbance of $NAD^+$ and NADH was measured using an analytical HPLC system described above (absorbance 340 nm). While $NAD^+$ is not fluorescent, NADH fluorescence was measured using its fluorescence wavelengths (ex = 340 nm, em = 460 nm).[8] The identity of $NAD^+$ and NADH was confirmed by comparing the retention time of samples to that of $NAD^+$ and NADH standards (Sigma Aldrich, 10127981001 and 10128023001, respectively). A calibration curve was performed using a formic acid standard (LC-MS grade, Fisher Chemical™, A117). Samples were prepared according to the kit's instructions and injected onto the analytical HPLC system equipped with a Poroshell column (4.6 × 150 mm, 2.7 μm particle size). The mobile phase contained 10 mM ammonium acetate (A) and acetonitrile (B). NADH signal was observed using a gradient of 5% to 30% acetonitrile (B) in ammonium acetate (A) over 50 min at 2.0 mL/min.

**α-MGH-1 and α-APY Antibody Screening for On-bead Peptide Glycation.** To test the specificity of the α-APY antibody, peptide 1 was synthesized using Fmoc-based solid phase peptide synthesis on ChemMatrix® resin (100-200 mesh, 0.51 mmol/g loading, PCAS BioMatrix, Inc., 1040378). On-bead peptide synthesis was adapted from protocols reported by McEwen et al.[4] After side chain deprotection, beads were washed with DMF (3 x 1 min), DCM (3 x 1 min), followed by ultrapure water (3 x 1 min), and allowed to equilibrate in ultrapure water overnight. The following day, on-bead peptides (2.55 μmol) were treated with 2 equiv. of MGO (5.1 μmol) in water and PBS (20 mM final concentration) and allowed to incubate at 37°C for 3 hours (or 24 hours followed by 24 hours of dilution for a total of 48 hours according to dilution protocols). Following incubation, beads were washed (3 x 1 min) with 20 mM Tris buffered with 150 mM NaCl containing 0.1% Tween (TBST) to remove excess MGO. Subsequently, beads were blocked using a 1% solution of bovine serum albumin (BSA) in TBST for 2 hours at room temperature. The blocked beads were then incubated with a 1:1000 dilution of either α-MGH-1 antibody (Cell BioLabs, Inc. STA-011) or α-APY antibody (Cosmo Bio Co., LTD., NOF-N213430-EX) for 18 to 20 hours at room temperature. After incubation with primary antibody, beads were washed with TBST (5 x 1 min) and exposed to a 1:1000 dilution of α-mouse secondary antibody (Abcam, AB7069) conjugated to alkaline phosphatase for 2 hours at room temperature. After incubation with secondary antibody, beads were washed with TBST (3 x 1 min) and alkaline phosphatase buffer (100 mM Tris-HCl, 150 mM NaCl, 1 mM $MgCl_2$ at pH 9.0) (3 x 1 min) and allowed to equilibrate in the buffer for 15 min. Afterwards, beads were exposed to color-developing reagents 5-bromo-4-chloro-3-indolyl phosphate and nitro blue tetrazolium (BCIP/NBT, Promega, S3771), washed with TBST (3 x 1 min), and transferred to a petri dish for microscopic imaging (Leica DMi8).

**Mammalian Cell Culture and Sample Preparation of TMT-labeled Peptides.** 150 mm culture dishes were seeded with 10 x $10^6$ HEK-293T cells in triplicates and grown to confluency in DMEM with 10% FBS supplementation. Cells were treated with 2 mM MGO in serum free DMEM for 24 hours and harvested using TrypLE Express (Gibco). Following MGO treatment, cell viability was accessed using a Trypan blue viability assay on a plate reader. Cell pellets were lysed in 200 μl ice cold 8 M Urea/50 mM triethylammonium bicarbonate (TEAB) with phosphatase/protease inhibitors (Pierce) and sonicated at 10% amplitude with 3 x 10 s pulses, followed by centrifugation at 15000 rpm for 15 min. Supernatant was collected, reduced with 8 mM dithiothreitol for 1 h, alkylated with 20 mM iodoacetamide for 30 min at room temperature in the dark. Subsequently, samples were digested overnight with trypsin at 37 °C. The digestion reaction was quenched using 0.1% trifluoroacetic acid, and samples were desalted using Pierce™ Peptide Desalting $C_{18}$ Spin Columns (#89852). Peptides were collected and dried using a speedvac prior to TMT labeling.

**TMT Labeling Protocol.** 100 μg of the dried peptide samples were reconstituted in 100 μL of 100 mM TEAB buffer. Each sample was then labeled using TMT 10-plex as per the manufacturer's instructions (TMT10plex™ Isobaric Label Reagents and Kits, #90111). The labeling reaction mixture was incubated at room temperature for 1 h and quenched using 5 μL of

5% hydroxylamine solution with incubation for 15 min at room temperature prior to pooling. The pooled sample was then dried, desalted and reconstituted in 0.1% formic acid for LC-MS/MS analysis. The 10 TMT channels are composed of three biological replicates (each with three technical replicates) and one pool sample, in which the sample ratio between the pool channel (i.e., channel 126) and channels 127N, 127C, 128N, 128C, 129N, 129C, 130N, 130C and 131 is expected to be 1. The experiment was carried out in two batches where the pooled samples (i.e. channel 126) served as the bridge channel. In the first batch, 3 biological replicates of the untreated samples were labeled with 127N, 127C, 128N TMT tags, 3 biological replicates of the samples treated with 0.5 mM MGO were labeled with 128C, 129N, 129C and 3 biological replicates of the samples treated with 1 mM MGO were labeled with 130N, 130C, 131 TMT tags. In the second batch, 3 biological replicates of the samples treated with 2 mM MGO were labeled with 127N, 127C, 128N TMT tags, 3 biological replicates of the untreated mitochondrial samples were labeled with 128C, 129N, 129C and 3 biological replicates of the mitochondrial samples treated with 1mM MGO were labeled with 130N, 130C, 131 TMT tags.

**Mass Spectrometry Data Acquisition for Proteomic Analysis.** LC-MS/MS data acquisition for TMT-labeled pooled sample was carried out using Thermo Scientific Orbitrap Exploris 240 (Thermo Fisher Scientific, Bremen, Germany) connected with Vanquish Neo UHPLC system (Thermo Fisher Scientific, Germany). Reconstituted peptides were loaded onto a trap column (300μm x 5mm, Thermo Scientific PepMap Neo Trap Cartridge, 174500). Peptides were separated on an analytical column (75μM x 150mm, Thermo Scientific EASY-Spray™ PepMap™ Neo UHPLC columns, ES75150PN) at a flow rate of 300 nL/min using an optimized linear gradient of 2%-45% acetonitrile for 235 min. Mass spectrometry data were acquired in data-dependent acquisition (DDA) mode with "top speed" of 3 sec cycle time. MS1 scans were acquired at a mass resolution of 60,000. The automatic gain control (AGC) target value for precursor ion acquisition was set to $1\times10^6$ and maximum ion injection time was set to automatic mode. Scan range of masses was at 350-1400 m/z. RF lens was set at 70%. Isolation width of 0.4 m/z was used for precursor ion selection and fragmented using higher-energy collision dissociation (HCD) with 36% normalized collision energy. MS2 scans were acquired at a mass resolution of 45,000 with AGC target as $1\times10^6$ and maximum ion injection time at auto mode. A FAIMS Pro Interface was connected to the Exploris 240 mass spectrometer. The compensation voltage (CV) of the FAIMS Pro interface was set to a combination of -45, -60 and -75 CVs. Each of the three CVs used was set to run DDA mode for 1 s cycle to build a big cycle of 3 s.

**Mass Spectrometry Proteomic Data Processing and Analysis.** All RAW files were processed and analyzed using Thermo Proteome Discoverer (PD) v.3.0. All database searches were performed using MSFragger-PD node (PMID: 36475762) as the combination of MSFragger and PeptideProphet (Philosopher) processing nodes. The default settings were predominantly used in MSFragger-PD node unless otherwise stated. Homo sapiens (SwissProt TaxID 9606, 20422 proteins) was used for the protein database. For all searches, trypsin cleavage was set to full, a precursor mass tolerance of 20 ppm was used, with a fragment mass tolerance of 0.6 Da allowing tryptic peptides only with up to two missed cleavages. The raw data was searched for five dynamic PTM combinations that could be found on peptides. The static modifications included carbamidomethylation (+57.021 Da) of cysteine and TMT-modification (+229.163 Da) of lysine and peptide N-terminus, and the dynamic modifications included argpyrimidine (+80.026 Da, R), methylglyoxal-derived hydroxyimidazolone isomers (MGH-isomers) (+54.011 Da, R), carboxyethyl arginine/lysine or MGH-DH (CEA/CEL/MGH-DH) (+72.021, R, K), tetrahydropyriminidine (THP) and/or other [M+144] AGEs (+144.041 Da, R), and phosphorylation (+79.966 Da; S, T, Y). A separate, parallel analysis was performed using the same static modifications but a different combination of dynamic modifications, including dimethylation (+28.031 Da, R), acetylation (+42.011 Da, R), argpyrimidine (+80.026 Da, R), and phosphorylation (+79.966 Da; S, T, Y). A "closed" search was performed to minimize the computational requirements for database searching for each technical replicate and all FAIMS parameters included in this study. PeptideProphet was used for PSM validation on these searches allowing a 1% FDR (false discovery rate). All bioinformatic analysis of LC-MS/MS data was performed in the R statistical computing environment. Gene Ontology (GO) enrichment analysis was performed using DAVID (PMID: 19131956) and proportional venn diagrams were generated using https://www.deepvenn.com/.

**MassIVE submission.** The mass spectrometry proteomics data have been deposited to the MassIVE repository with the dataset identifier MSV000097149.

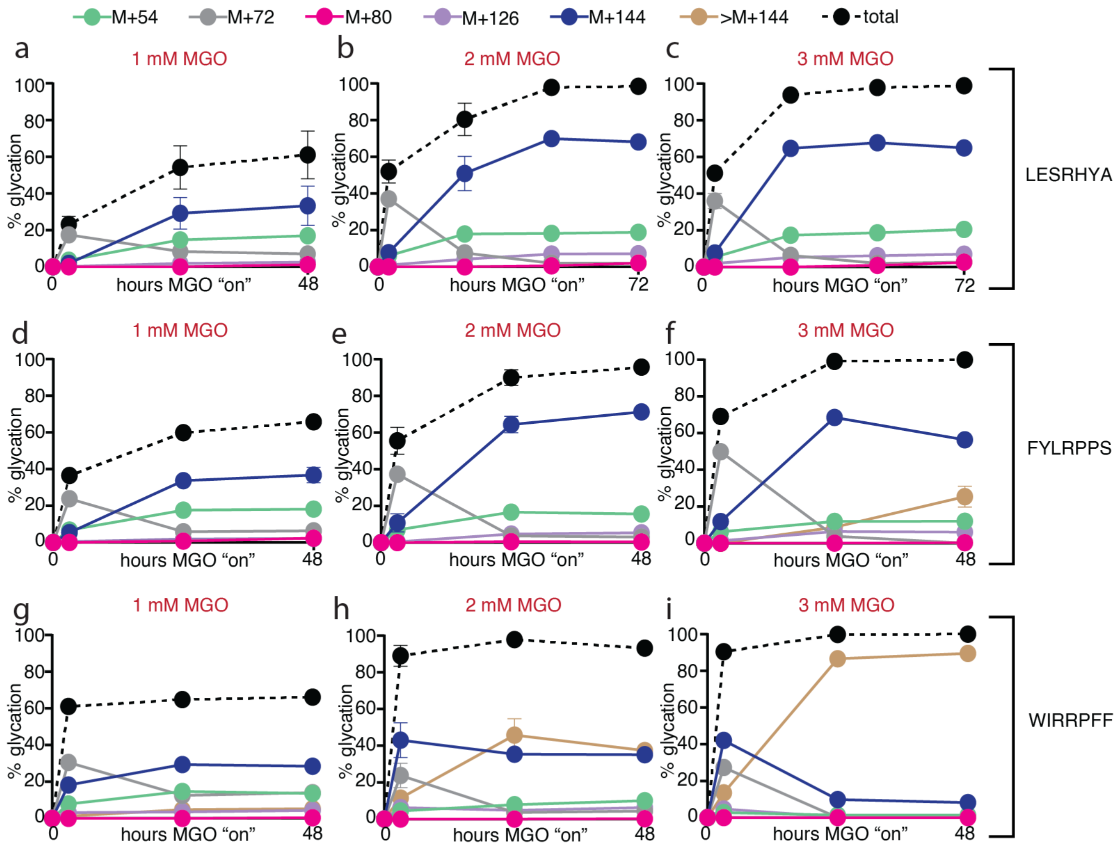


**Figure S1. Peptide Glycation for Previously-Identified APY-modified Protein Sequences. (a), (b) & (c)** Distribution of AGEs on MGO-treated peptide **1** at 1 mM, 2 mM, and 3 mM MGO concentrations after 3 h, 24 h, 48 h, or 72 h ($n \geq 3$). **(d), (e) & (f)** Distribution of AGEs on MGO-treated peptide Ac-FYLRPPS at 1 mM, 2 mM, and 3 mM MGO concentrations after 3 h, 24 h, and 48 h ($n \geq 3$). **(g), (h) & (i)** Distribution of AGEs on MGO-treated peptide Ac-WIRRPFF at 1 mM, 2 mM, and 3 mM MGO concentrations after 3 h, 24 h, and 48 h ($n = 2$ for 48 h time point, $n = 3$ for all other time points). These peptides were chosen because they have been identified as APY-modified based on previous work done in our lab.[4,9] Specifically, peptide Ac-LESRHYA was identified via a one-bead-one-compound combinatorial peptide library and selected for its ability to form MGH-1. Peptides Ac-FYLRPPS and Ac-WIRRPFF, both from lens crystallin, were identified by in vitro glycation of proteins. In these studies, all three peptides were identified with APY formation upon treatment with MGO. Though APY formation requires two MGO, increasing initial MGO concentrations did not increase APY levels even after 48 hours of incubation.

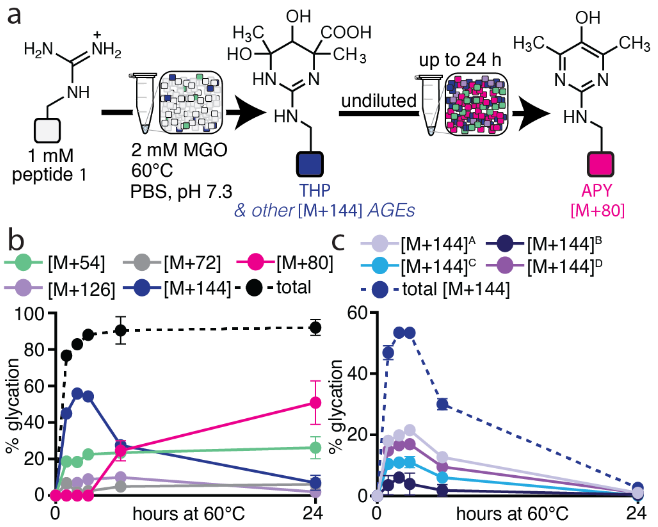


**Figure S2. All [M+144] Species Mature into APY over Time.** General scheme describing the experimental procedure of glycation reactions on MGO-treated peptide **1** 60 °C (20 mM PBS, pH 7.3). Time points were taken at 0 h, 1 h, 2 h, 3 h, 6 h, and 24 h to monitor changes in AGE distributions (n ≥ 3). **(b)** Distribution of AGEs on MGO-treated peptide **1**, also shown in **Main Text Fig. 3**. **(c)** Distribution of all [M+144] AGEs observed on MGO-treated peptide **1**. We consistently observed up to four different [M+144] species with distinct retention times. To our knowledge, THP is the only [M+144] AGE that has been reported.[10] However, all of the [M+144] species we observed appear to mature into APY, since each one decreased as APY levels increased. Therefore, we report [M+144] as a total sum of all [M+144] AGEs.

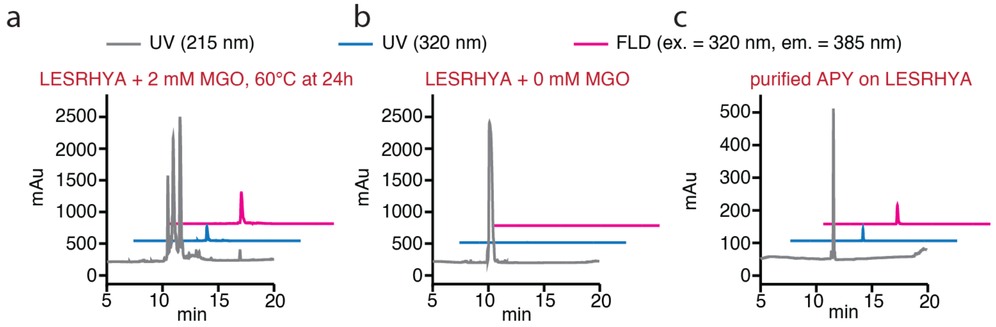


**Figure S3. Detection of APY Fluorescence by Analytical HPLC.** The characteristic fluorescence wavelengths of APY (ex = 320 nm, em = 385 nm) have been reported and can be measured using an HPLC system equipped with a fluorescence detector.[1–3] (**a**) HPLC chromatogram for the reaction of peptide **1** treated with MGO at 60 °C. (**b**) HPLC chromatogram for peptide **1** in the absence of MGO at 0 h. (**c**) HPLC chromatogram for purified APY-modified peptide **1** (peptide $\mathbf{1}^{\mathbf{APY}}$). Absorbance at 215 nm and 320 nm was collected to observe peptide bond and APY absorbance wavelengths, respectively. Fluorescence excitation at 320 nm and emission at 385 nm were collected based on reported APY characteristic fluorescence wavelengths. APY absorbance and fluorescence peaks were observed in both the reaction and purified peptide $\mathbf{1}^{\mathbf{APY}}$ samples in (**a**) & (**c**), yet were absent in the control sample (**b**), confirming the presence and identity of APY in our samples.

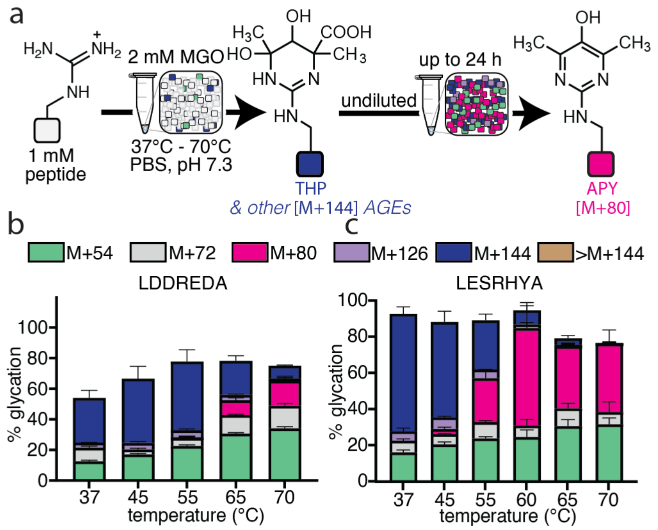


**Figure S4. APY Formation can be Affected by the Surrounding Sequence.** (**a**) General scheme describing the temperature scan conducted for 24 h peptide glycation reactions performed at 37 °C, 45 °C, 55 °C, 65 °C, or 70 °C (n ≥ 3 for all reactions). (**b**) Distribution of AGEs on MGO-treated peptide LDDREDA after 24 h. (**c**) Distribution of AGEs on MGO-treated peptide Ac-LESRHYA after 24 h, also shown in **Main Text Fig. 3**. Ac-LDDREDA was identified as a negative hit sequence from a previous combinatorial peptide library for MGH-1 formation and was not known to form APY.[4] Since clustered negative charges have been reported to impede glycation, it was important to confirm that this also included APY formation. Expectedly, overall glycation levels were seen to decrease for Ac-LDDREDA compared peptide **1**. Although APY levels increased as temperature increased for both sequences, we found that APY was not formed until higher temperatures for Ac-LDDREDA.

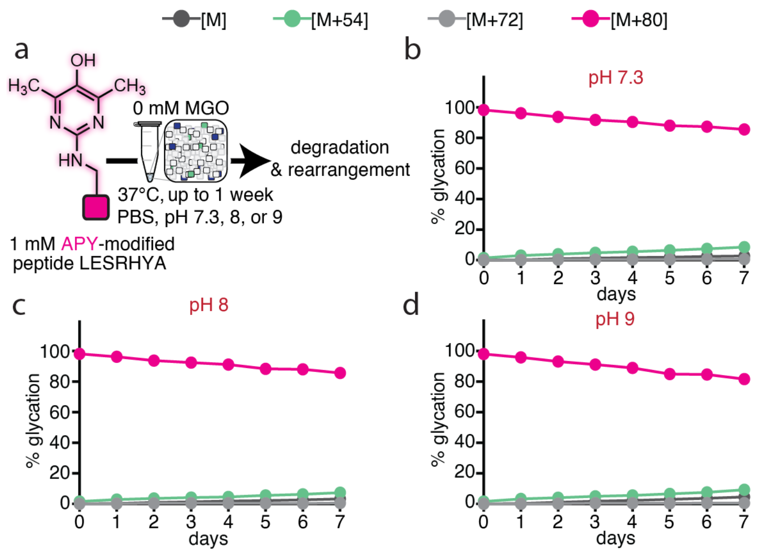


**Figure S5. APY on a Short Peptide is Stable under Physiological Conditions**. APY has been reported to be unstable at pH 7.4 and 37 °C, having a short lifetime (1.7 days), which was extended to 9.3 days in the presence of the metal-ion chelator DETAPAC.[11] However, our extensive peptide studies using peptide **1** suggested that APY is more stable than previously reported. We purified APY-modified peptide **1** (peptide $\mathbf{1}^{\mathbf{APY}}$) to conduct stability studies in PBS with pH 7.3, 8, and 9 ($n \geq 3$ for all reactions). We found that APY was much more stable than previously thought, exhibiting a steady but slow decay even after 7 days of incubation in PBS pH 7.3 at 37 °C, without any metal-ion chelator. As pH increased, APY decayed faster, and a mass adduct [M+54] with the same retention time as that of MGH-1 was observed. The difference in results of APY stability could be due to the effects of the microenvironment surrounding the APY site on a short peptide, which was not present in previous studies using monomeric APY. Together, our results confirmed that APY—or AGE overall—formation and stability are greatly influenced by the surrounding amino acids of the glycated site, thereby suggesting that glycation studies done on peptides or proteins can offer different insights into the behaviors of AGEs that are more likely to be observed in a cellular context.

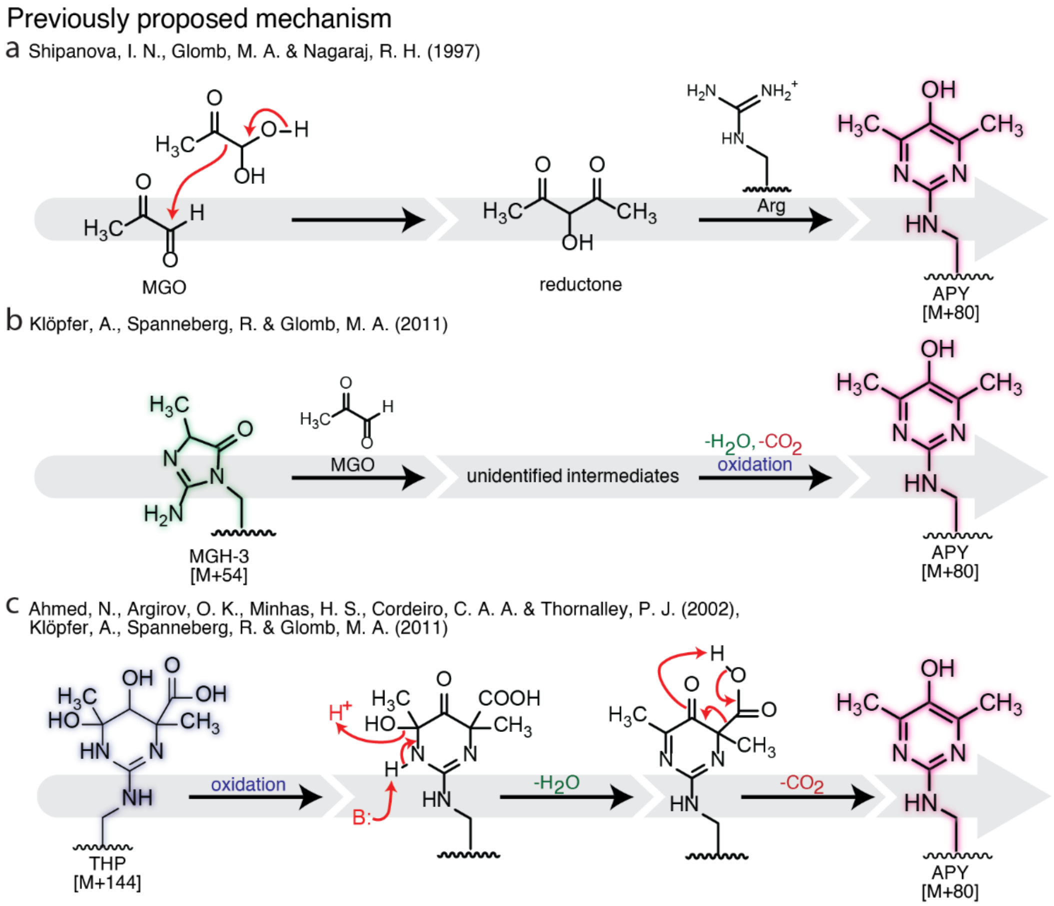


**Figure S6. Previously Proposed APY Formation Mechanisms.** Previously proposed mechanisms for APY formation from (**a**) reductone formation, (**b**) MGH-3 via oxidative decarboxylation, and (**c**) THP via oxidative decarboxylation.[1,2,11]

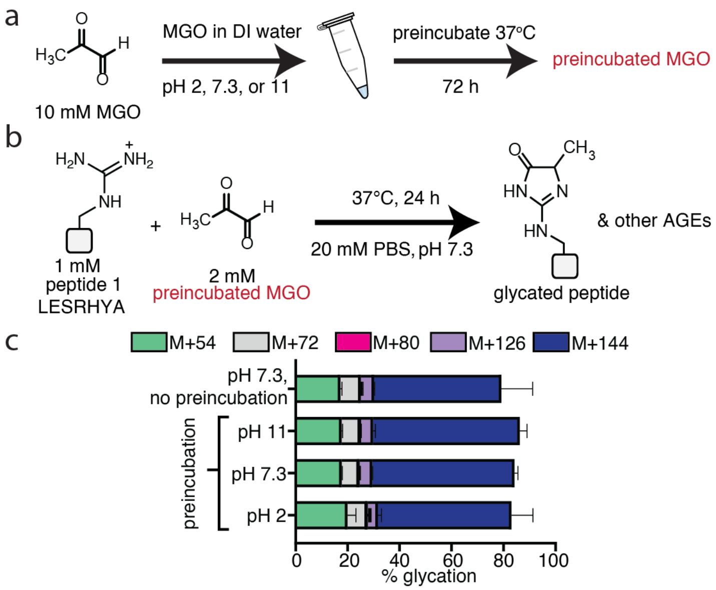


**Figure S7. MGO Preincubation at Different pH does not generate APY.** APY has been proposed to form from the reaction between Arg and a reductone intermediate, which results from the reaction between two MGO (see **Figure S6**).[2] Thus, we preincubated MGO to see if this would facilitate APY formation by generating reductone prior to reaction with peptide. General scheme describing (**a**) the preincubation of MGO in ultrapure water at 37 °C, with pH adjusted to 2, 7.3, or 11, and (**b**) peptide glycation reactions using preincubated MGO. (**c**) Distribution of AGEs on MGO-treated peptide **1** after 24 h of incubation (n = 3). A control sample, labeled "no preincubation", was also performed for comparison. We found virtually no difference in not only APY levels but also other AGE distributions, indicating that APY is unlikely to be formed *via* the reaction between Arg and an MGO reductone.

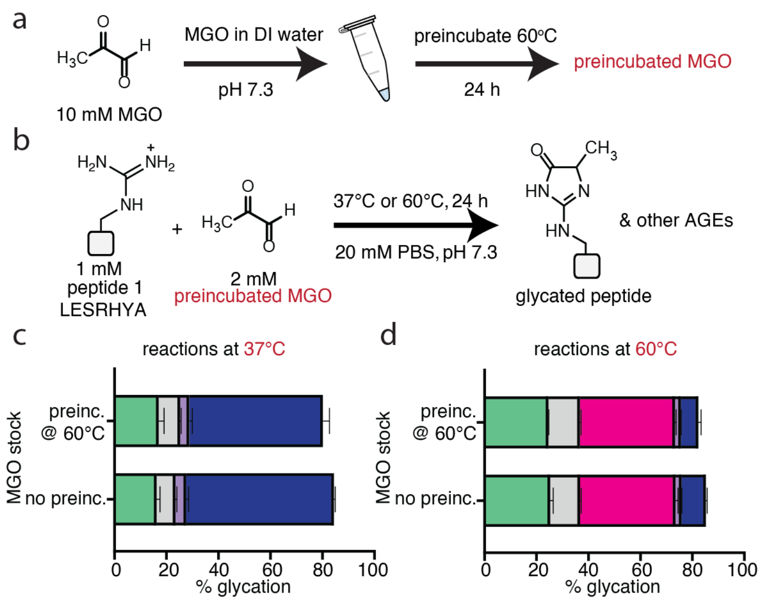


**Figure S8. MGO Preincubation at Elevated Temperatures does not produce APY.** As the formation of the previously proposed reductone intermediate could also be dependent on temperature,[2] we preincubated MGO at an elevated temperature. General scheme describing **(a)** the preincubation of MGO in ultrapure water at 60 °C, in neutral pH and **(b)** peptide glycation reactions using preincubated MGO. **(c)** Distribution of AGEs on MGO-treated peptide **1** after 24 h of incubation at 37 °C (n = 3). A control sample, labeled "no preinc.", was also performed for comparison. There was virtually no difference in AGE distributions between these samples. **(d)** Distribution of AGEs on MGO-treated peptide **1** after 24 h of incubation at 60 °C (n = 3). A control sample, labelled "no preinc.", was also performed for comparison. No differences were observed for AGE distributions and APY levels between these samples, providing further evidence that the reductone mechanism is unlikely.

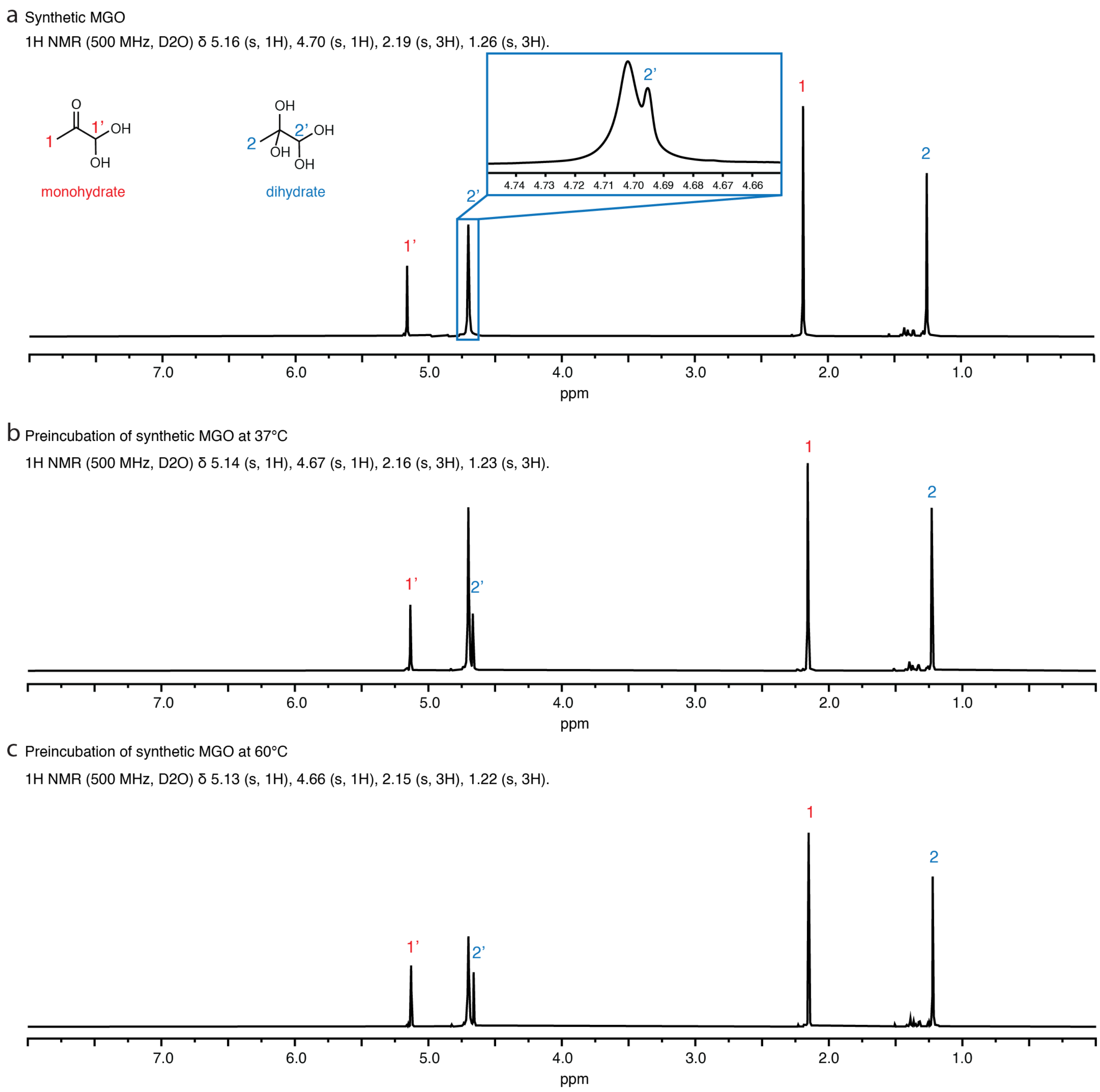


**Figure S9. NMR Spectra do not show Reductone Formation by MGO Preincubation.** (**a**) NMR spectrum of a fresh stock of synthetic MGO. (**b**) NMR spectrum of synthetic MGO following preincubation at 37 °C for 72 h. (**c**) NMR spectrum of synthetic MGO following preincubation protocol at 60 °C for 72 h. The peak with a shift of 4.7 corresponds to the solvent peak, sometimes not entirely resolvable with the aldehyde proton (labeled as "**2'**") peak of the dihydrate form. Monohydrate (red) and dihydrate (blue) forms of MGO were observed. A cluster of baseline peaks around 1.5 ppm region have been reported as a result of MGO polymerization.[12] Importantly, in both forms, the integration of the aldehyde proton (1H) peak (**1'** or **2'**) is one-third of that of the methyl proton (3H) peak (**1**, **2**). This would not be the case for in the MGO reductone. The NMR spectra were observed to be identical before and after incubation, indicating no reductone formation.

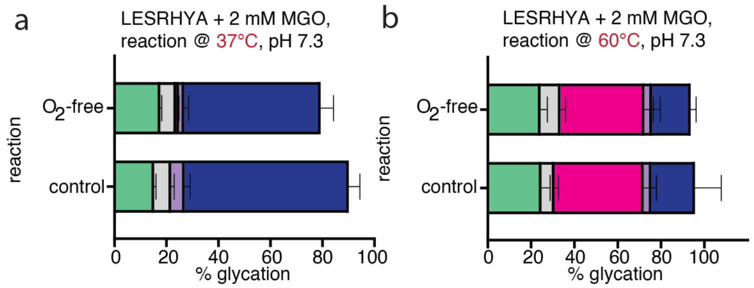


**Figure S10. Oxygen is Not Required to Form APY.** Previous studies suggested that APY was formed from THP rearrangement *via* a process involving an undetermined oxidation step.[1,11] To investigate the role of oxygen in APY formation from THP and other [M+144] AGEs, we conducted glycation reactions with peptide **1** in the presence (control) or absence ($O_2$-free) of oxygen. All reactions components were carefully degassed with $N_2$, and reactions were assembled inside a glovebox filled with $N_2$ ($O_2$<4ppm). Control samples followed our standard glycation protocols, without any degassing steps. (**a**) Distribution of AGEs on MGO-treated peptide **1** after 24 h of incubation at 37 °C (n = 3). Despite a slightly lowered overall glycation, AGE distributions remained unchanged when comparing $O_2$-free samples to control samples. (**b**) Distribution of AGEs on MGO-treated peptide **1** after 24 h of incubation at 60 °C (n = 3), showing virtually no difference in APY levels observed between samples. Together, these data showed that APY formation from [M+144] rearrangement was not dependent on the presence of oxygen, suggesting that an alternative mechanism does not require a formal oxidation step.

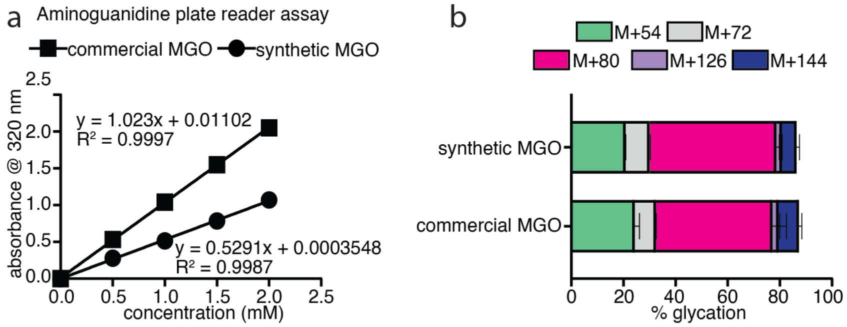


**Figure S11. Synthetic and Commercial MGO Sources both generate APY.** As our glycation reactions used commercial MGO, which may contain impurities, we aimed to investigate whether APY formation was affected by the use of commercial MGO compared to a freshly synthesized stock of MGO. (**a**) Following the successful synthesis of MGO, an aminoguanidine assay was performed to correct for any differences in MGO concentrations between the two sources using the slope of the calibration curve. (**b**) Distribution of AGEs on MGO-treated peptide **1** after 24 h of incubation at 60 °C (n = 3). These results showed no differences in APY levels or AGE distributions between samples, confirming that APY formation remained unaffected by either source of MGO used.

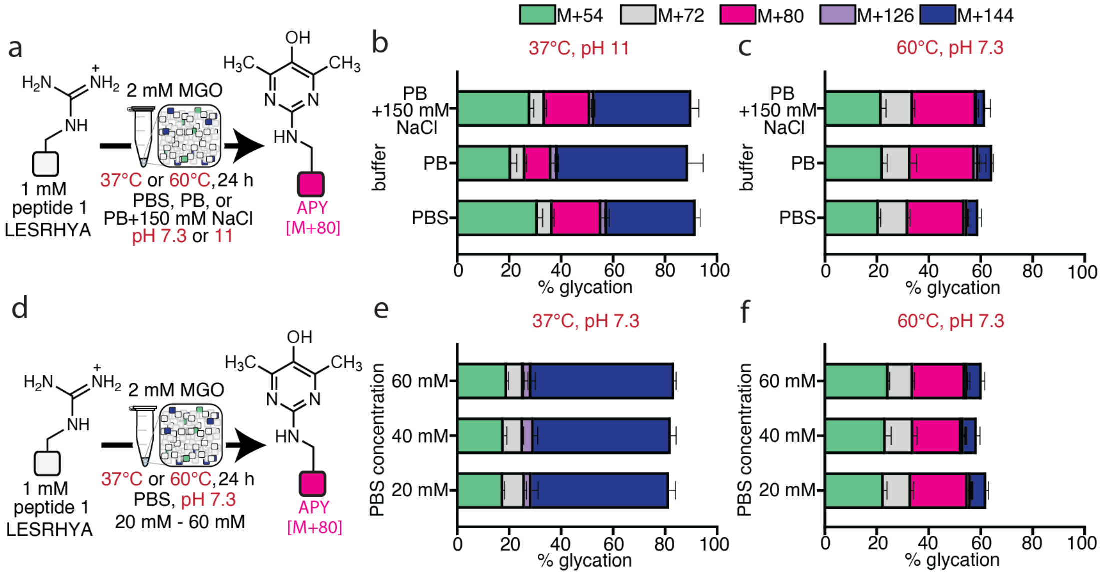


**Figure S12. Phosphate Buffer and Salt Concentrations do not impact APY formation.** To evaluate how phosphate concentration and buffer salts affected APY formation, we performed a series of controls using peptide **1**. **(a)** General scheme describing glycation reactions of peptide **1** using differently prepared buffers (24 h incubation, 37 °C or 60 °C, pH 7.3). Buffers used included a commercial phosphate buffered saline (PBS, which contained 100 mM sodium phosphate and 150 mM sodium chloride), phosphate buffer (PB) which was prepared in-house by adding calculated amounts of monobasic monohydrate and dibasic sodium phosphate in ultrapure water, and phosphate buffer with 150 mM NaCl (PB + 150 mM NaCl). **(b)** Distribution of AGEs on MGO-treated peptide **1** after 24 h of incubation at 37 °C (n = 3). **(c)** Distribution of AGEs on MGO-treated peptide **1** after 24 h of incubation at 60 °C (n = 3). **(d)** General scheme describing glycation of peptide **1** at standard reaction conditions using PBS at varied concentrations of 20 mM, 40 mM, and 60 mM. **(e)** Distribution of AGEs on MGO-treated peptide **1** after 24 h of incubation at 37 °C (n = 3). **(c)** Distribution of AGEs on MGO-treated peptide **1** after 24 h of incubation at 60 °C (n = 3). These results demonstrate that salt and buffer concentration had hardly any effects on APY formation.

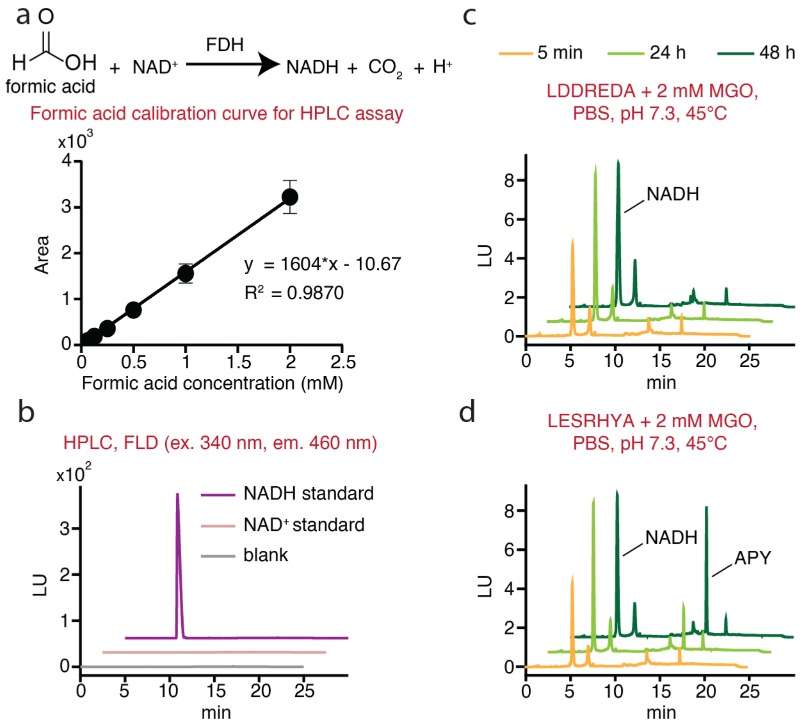


**Figure S13. Detection of Formic Acid during Glycation Reactions.** Our proposed mechanism shows formic acid as a byproduct of [M+144] rearrangement. Therefore, we aimed to identify formic acid using a formic acid assay kit purchased from Megazyme. Briefly, formate dehydrogenase (FDH) was used to detect formic acid in samples by converting $NAD^+$ to NADH, the fluorescence wavelengths of which was measured using HPLC equipped with a fluorescence detector (ex = 340 nm, em = 460 nm).[8] **(a)** General scheme depicting the assay design, and the calibration curve obtained using a formic acid standard (n = 3). **(b)** HPLC chromatogram of the working assay using NADH and $NAD^+$ standards, showing NADH fluorescence signal (ex = 340 nm, em = 460 nm) that was not observed in the $NAD^+$ sample. The retention time of NADH was observed around 5 min. **(c) & (d)** HPLC chromatograms of MGO-treated peptide LDDREDA (upper right) and peptide **1** (lower right) under standard glycation reaction conditions (24 h incubation, 45 °C, pH 7.3). 45 °C was chosen as the reaction temperature because at this temperature, we expected peptide **1** to produce observable APY signals, hence NADH signals, while this would not be the case for peptide LDDREDA. While higher concentration of formic acid was detected in 24 h reactions of peptide **1** with 2 mM MGO, similar concentrations of formic acid were observed when compared between peptide **1** and LDDREDA which had significantly less APY under the same condition. These results indicate that while formic acid may be a byproduct of our proposed mechanism, it is likely not exclusive to APY formation.

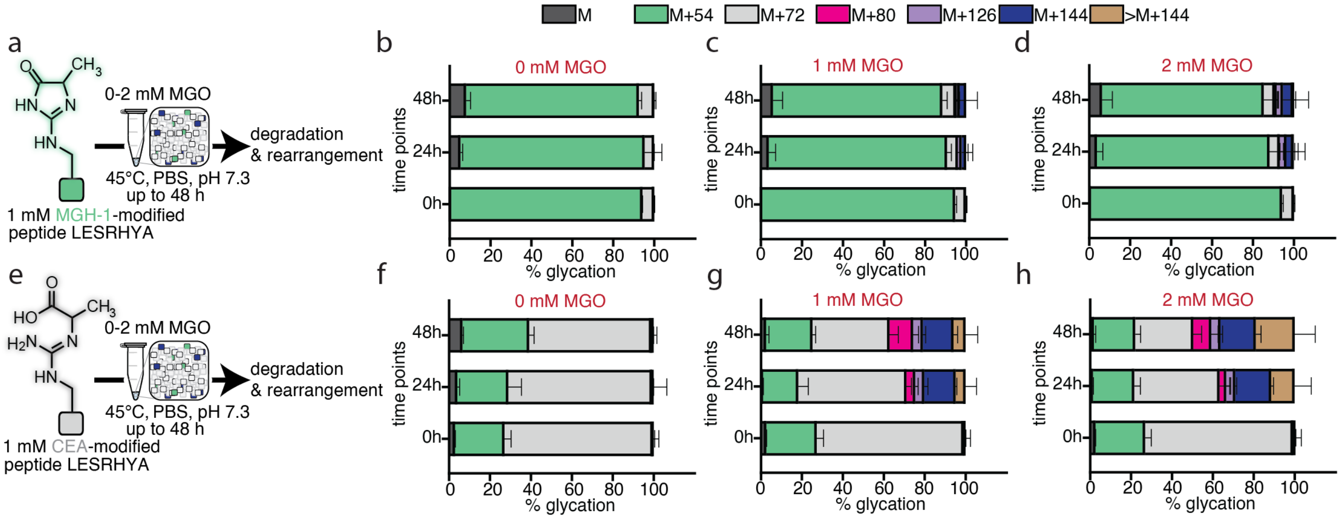


**Figure S14. CEA is also a Precursor for APY.** Just like THP, APY formation requires two moieties of MGO to react with an Arg side chain.[2,10,13] In our proposed mechanism, we noticed an intermediate structure that resembles a tautomer of CEA, allowing us to hypothesize that CEA might be a precursor to APY. In order to test this idea, we reacted peptide **1** to form CEA ($\mathbf{1}^{\mathbf{CEA}}$) and MGH-1 ($\mathbf{1}^{\mathbf{MGH\text{-}1}}$) according to previously reported protocols and purify these AGE-modified peptides. Due to coelution, peptide $\mathbf{1}^{\mathbf{CEA}}$ contained 20% MGH-1 and 80% CEA. **(a) & (e)** General schemes describing the experimental procedure of glycation reactions on MGO-treated peptide $\mathbf{1}^{\mathbf{MGH\text{-}1}}$ (upper left) and peptide $\mathbf{1}^{\mathbf{CEA}}$ (lower left) at 0 h, 24 h, and 48 h of incubation (45 °C, pH 7.3). The reaction temperature was set 45 °C to expedite APY formation. **(b) & (f)** Distributions of AGEs on peptide $\mathbf{1}^{\mathbf{MGH\text{-}1}}$ (upper) and peptide $\mathbf{1}^{\mathbf{CEA}}$ (lower) without MGO (0 mM) after 24 h of incubation at 45 °C (n = 3). **(c) & (g)** Distributions of AGEs on peptide $\mathbf{1}^{\mathbf{MGH\text{-}1}}$ (upper) and peptide $\mathbf{1}^{\mathbf{CEA}}$ (lower) treated with 1 mM MGO after 24 h of incubation at 45 °C (n = 3). **(d) & (h)** Distributions of AGEs on peptide $\mathbf{1}^{\mathbf{MGH\text{-}1}}$ (upper) and peptide $\mathbf{1}^{\mathbf{CEA}}$ (lower) treated with 2 mM MGO after 24 h of incubation at 45 °C (n = 3). Upon treatment with MGO, only peptide $\mathbf{1}^{\mathbf{CEA}}$ was observed with APY at 24 h and 48 h, confirming our hypothesis that CEA is another potential precursor for APY.

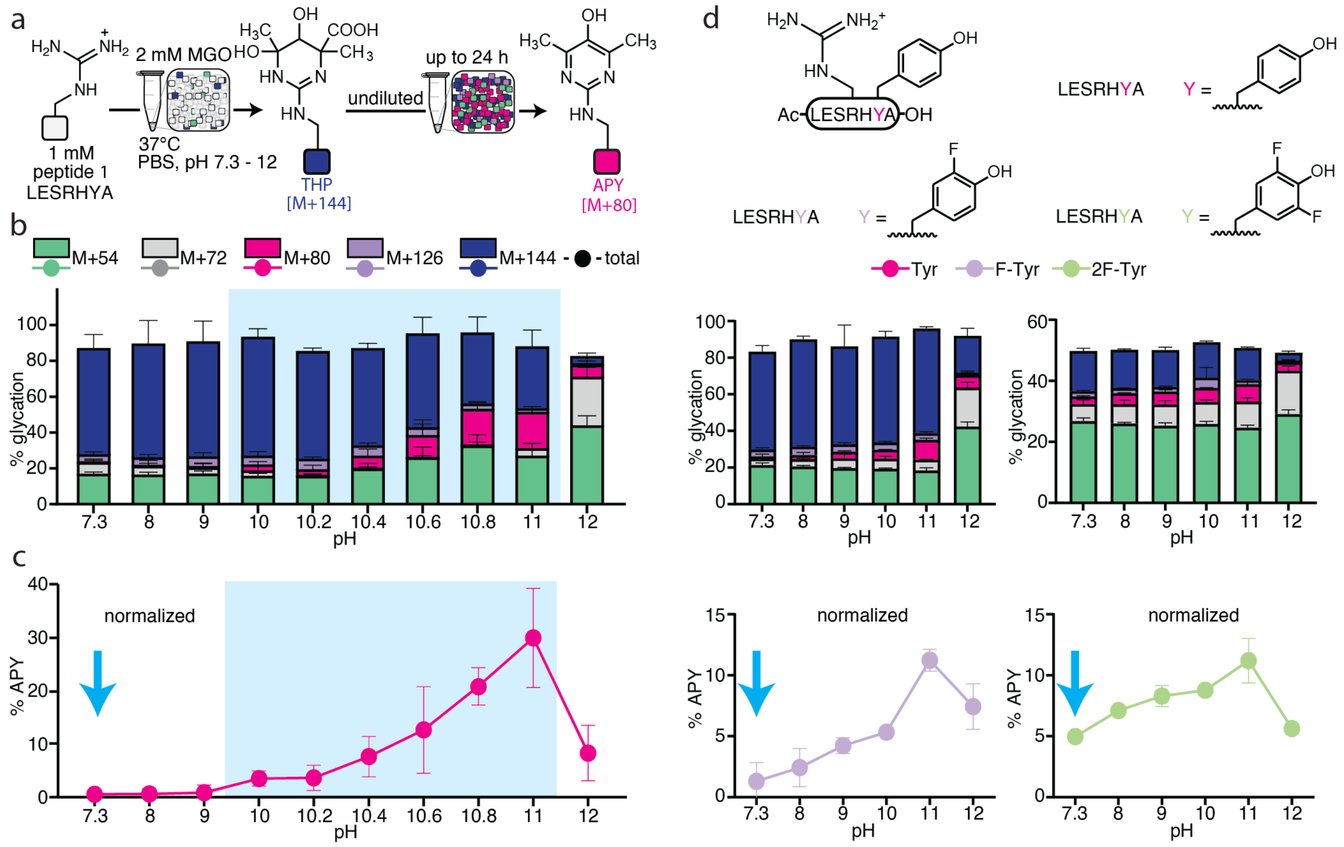


**Figure S15. MGO Treatments of Fluorinated Variants of Peptide 1.** To test the hypothesis that a nearby Tyr facilitates APY formation, we synthesized variants of peptide **1** that incorporated noncanonical Tyr derivatives with lowered pKa values. Specifically, 3-fluorotyrosine (peptide **1**$^{F}$) and 3,5-difluorotyrosine (peptide **1**$^{FF}$) were chosen because of their pKa values (8.4 and 7.2, respectively) in comparison with the pKa of Tyr (10.3).[14] **(a)** General scheme describing the experimental procedure of a pH screening of glycation reactions on MGO-treated peptide **1** using PBS buffer with pH in a range of 7.3—12 (37 °C, 24 h incubation). **(b)** Distribution of AGEs on MGO-treated peptide **1** after 24 h of incubation at 37 °C (pH 7.3—12). A short pH scan (in the blue box) was also performed for reactions between pH 10—11 in increments of 0.2 (n≥3 for all reactions). **(c)** % APY was normalized to % total glycation. We observed a sharp transition in APY levels between pH 10.2 and 10.4. This pH range corresponds to the reported pKa value for Tyr (10.3), suggesting Tyr plays an active role in APY formation. **(d)** Distributions of AGEs on MGO-treated peptide **1**$^{F}$ and peptide **1**$^{FF}$ after 24 h of incubation at 37 °C (pH 7.3—12) (center) and % APY for these reactions when normalized to % total glycation (lower). Overall glycation was seen to decrease as more F was substituted on Tyr, which was expected as clustered negative charges impeded glycation. For reactions with peptide **1**$^{FF}$, MGH-1 was now the predominant AGE instead of [M+144]. This observation was also consistent with previous work done in our lab on chlorinated peptide **1**.[4] The distribution of APY across the pH range between 7.3 and 12 exhibited the same trend, with the highest amount of [M+80] observed at pH 11 for all three peptides. Between pH 7.3 and 10, % APY (both absolute and normalized) in peptide **1**$^{FF}$ was the highest, followed by peptide **1**$^{F}$, then peptide **1**. These results demonstrated the importance of the nearby Tyr as a facilitator for [M+144] rearrangement into APY. This matches our hypothesis that the presence of a nearby Tyr can fulfill the role of a general base, facilitating APY formation according to out proposed mechanism.

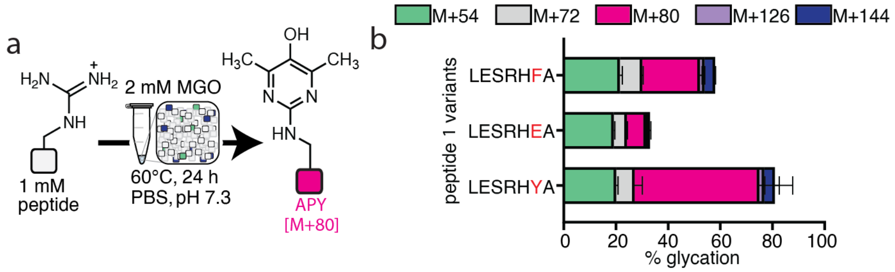


**Figure S16. Replacing the Tyr with Glu and Phe on Peptide 1 reduced Glycation and APY Formation.** In order to further confirm the role of Tyr as a general base that can facilitate APY formation on peptide **1**, we synthesized two other variants of peptide **1** that incorporated a negatively charged residue Glu (peptide $\mathbf{1}^{\mathbf{Glu}}$) and a non-polar residue Phe (peptide $\mathbf{1}^{\mathbf{Phe}}$) in place of Tyr. (**a**) General scheme describing the experimental procedure of glycation reactions on MGO-treated peptide **1**, $\mathbf{1}^{\mathbf{Glu}}$, and $\mathbf{1}^{\mathbf{Phe}}$ at 60 °C (pH 7.3, 24 h incubation) (n = 3). (**b**) Distribution of AGEs on MGO-treated peptide **1**, $\mathbf{1}^{\mathbf{Glu}}$, and $\mathbf{1}^{\mathbf{Phe}}$. Upon treatment with 2 mM MGO at 60 °C for 24 h, both peptide $\mathbf{1}^{\mathbf{Glu}}$ and $\mathbf{1}^{\mathbf{Phe}}$ expectedly exhibited less overall glycation and APY formation than peptide **1**, confirming the importance of Tyr on peptide **1** as a facilitator for APY formation.

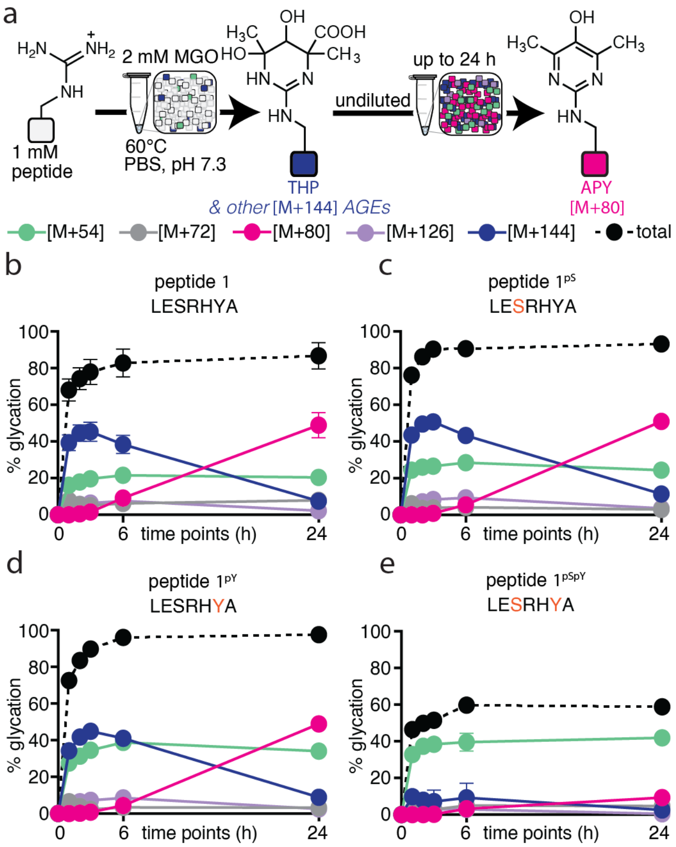


**Figure S17. Phosphorylated Amino Acids bias Certain AGE Formation at Elevated Temperatures.** To investigate the effect of phosphorylation on APY formation, we synthesized peptide **1**$^{pS}$ (a variant of peptide **1** containing pSer in lieu of Ser), peptide **1**$^{pY}$ (pTyr in lieu of Tyr), and peptide **1**$^{pSpY}$ (containing both pSer and pTyr). (**a**) General scheme describing the experimental procedure of glycation reactions on MGO-treated peptide **1**, **1**$^{pS}$, **1**$^{pY}$, and **1**$^{pSpY}$ at 60 °C (pH 7.3, 24 h incubation). Time points were taken at 0 h, 1 h, 2 h, 3 h, 6 h, and 24 h to monitor changes in AGE distributions. (**b**) Distribution of AGEs on MGO-treated peptide **1**. (**c**) Distribution of AGEs on MGO-treated peptide **1**$^{pS}$. (**d**) Distribution of AGEs on MGO-treated peptide **1**$^{pY}$. (**e**) Distribution of AGEs on MGO-treated peptide **1**$^{pSpY}$. (n = 3 for all reactions). There was no difference in AGE distributions and no noticeable difference in APY levels between peptide **1**, **1**$^{pS}$, and **1**$^{pY}$, despite the introduction of up to two negative charges. However, MGH-1 levels of peptide **1**$^{pS}$ and **1**$^{pY}$ were slightly higher than those of peptide **1**. This observation is consistent with our previous report on MGH-1 formation, which can be facilitated by a nearby Tyr residue acting as a general base. In the case of peptide **1**$^{pS}$, both pSer and Tyr can fulfill this role. When introducing up to four negative charges on peptide **1**$^{pSpY}$, overall glycation and APY levels decreased, but MGH-1 levels remained as the predominant adduct across all time points. These observations also support our proposal that a general base, such as Tyr or pSer, can be critical for APY formation, counteracting the hampering effects of negative charges on glycation.

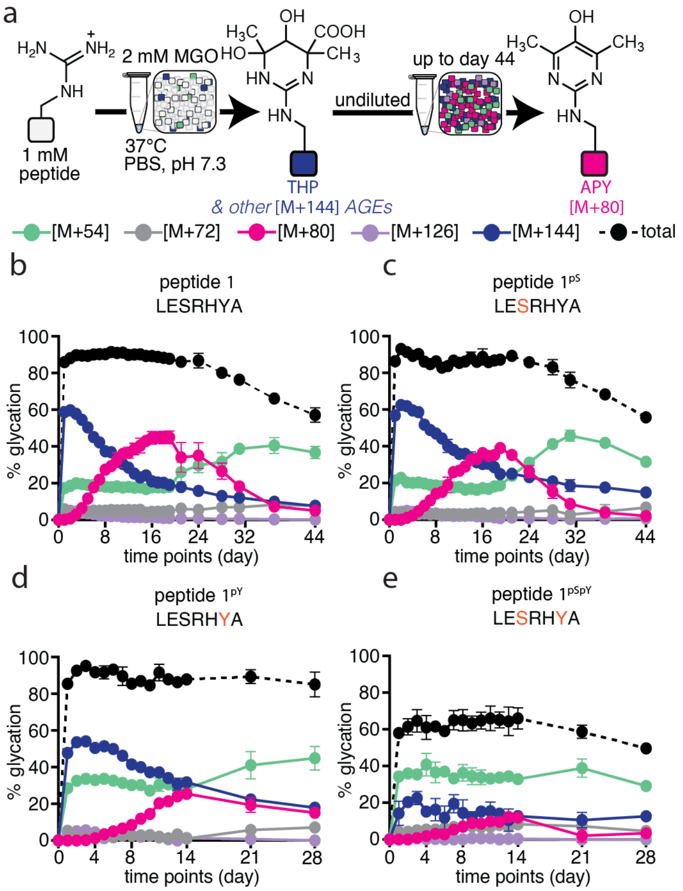


**Figure S18. Phosphorylated Amino Acids bias Certain AGE Formation at Physiological Temperature.** We extended our glycation experiments with peptide **1**, **1**$^{pS}$, **1**$^{pY}$, and **1**$^{pSpY}$ described in **Main Text Fig. 5** and **Supplementary Fig. S17** up to 44 days of incubation for peptide **1** and **1**$^{pS}$, 28 days for peptide **1**$^{pY}$ and peptide **1**$^{pSpY}$ at 37 °C. (**a**) General scheme describing the experimental procedure of glycation reactions on MGO-treated peptides at 37 °C (pH 7.3). Time points were taken every 24 h for the first 14 days (19 days for peptide **1** and **1**$^{pS}$), then on day 21, 24, and 28 (and on day 31, 37, and 44 for peptide **1** and **1**$^{pS}$). (**b**) Distribution of AGEs on MGO-treated peptide **1**. (**c**) Distribution of AGEs on MGO-treated peptide **1**$^{pS}$. (**d**) Distribution of AGEs on MGO-treated peptide **1**$^{pY}$. (**e**) Distribution of AGEs on MGO-treated peptide **1**$^{pSpY}$ (n = 2 for peptide **1**$^{pS}$ reactions collected on day 9 & 31; n = 2 for peptide **1** reactions collected on day 37; n = 2 for peptide **1**$^{pSpY}$ reactions collected on day 6; n = 3 for all other reactions). There was no difference in AGE distributions and no noticeable difference in APY levels between peptide **1** and **1**$^{pS}$, despite the introduction of up to two negative charges. We found that APY levels for both peptides peaked on day 19 and gradually decreased for the remaining time in this study. Interestingly, MGH-1 levels remained mostly unchanged in the first 15 days, then gradually increased and peaked on day 31, and decayed steadily for the remaining time in this study. These observations are consistent with previous reports, citing that MGH-1 is a stable AGE.[1,15] As MGH-1 levels started to increase on day 16, we noticed that APY levels also began to decrease and [M+144] continued to decrease, suggesting a mechanistic correlation between these three AGEs. While MGH-1 levels significantly increased for peptide **1**$^{pY}$ and peptide **1**$^{pSpY}$ at earlier time points and remained fairly constant throughout the incubation period, MGH-1 was the predominant AGE at all time points for only peptide **1**$^{pSpY}$ despite the lowered overall glycation due to four negative charges, suggesting that for peptide **1**$^{pSpY}$, phosphorylated residues bias MGH-1 formation over APY formation. Finally, we also noticed a continuous decay of an unknown adduct [M+126], though this adduct was observed consistently in low quantities (< 5%). In our proposed mechanism, this mass change potentially corresponds to an intermediate undergoing the final retro-aldol step to release formic acid and generate APY.

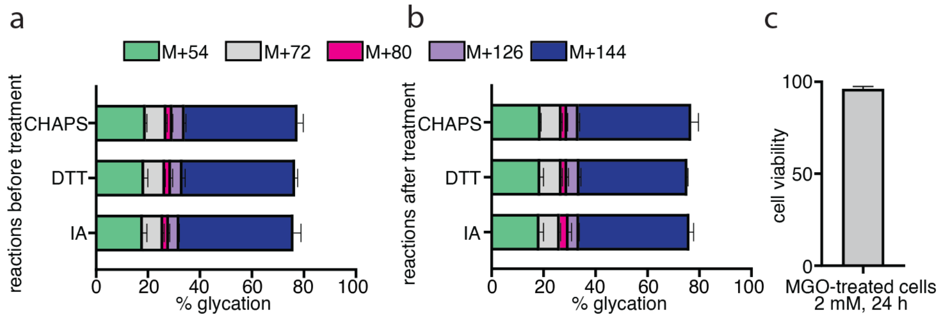


**Figure S19. Cell Viability and Controls showing that Proteomic Reagents do not impact the AGE Distribution.** Sample preparation for proteomics regularly involves reagents such as (3-((3-cholamidopropyl) dimethylammonio)-1-propanesulfonate) (CHAPS), dithiothreitol (DTT), and iodoacetamide (IA). To ensure that these reagents do not affect APY formation, we followed standard glycation reaction protocols on MGO-treated peptide **1** (2 mM MGO, pH 7.3, 37 °C, 24 h incubation), then treated AGE-modified peptide **1** with each reagent in a similar manner to how the reagent would be used as described in our proteomic protocols. Distributions of AGEs were compared before and after treatment of each reagent. (**a**) Distribution of AGEs on MGO-treated peptide **1** before treatment with proteomic reagents. (**b**) Distribution of AGEs on MGO-treated peptide **1** after treatment with proteomic reagents (n = 3 for all reactions). These results confirmed that proteomic reagents used in our studies would neither interfere with AGE distributions nor APY formation. (c) Cell viability of MGO-treated HEK-293T cells was measured post 24 h treatment using a Trypan blue viability assay on a plate reader (n = 3). More than 95% of treated cells remained viable, confirming that MGO treatment did not significantly decrease cell viability.

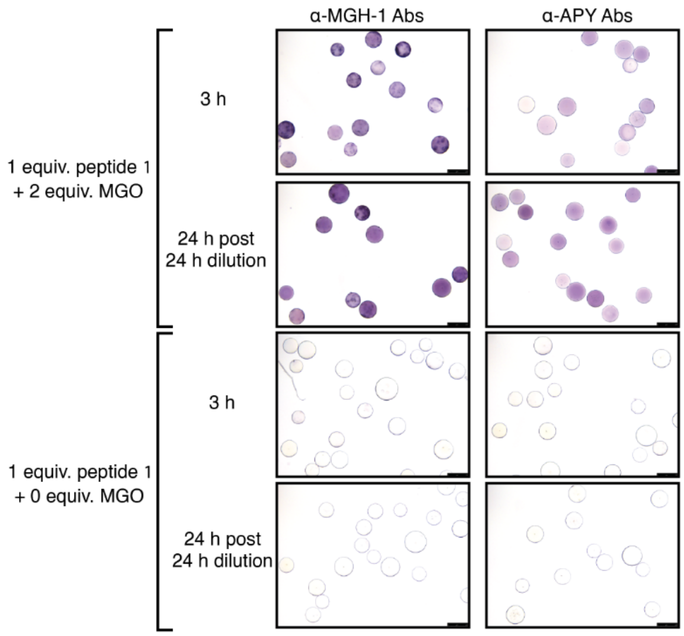


**Figure S20. α-MGH-1 and α-APY Antibody Screening for On-bead Peptide Glycation.** Upon treating on-bead peptide **1** with 2 equiv. of MGO, we observed dark colored purple beads when beads were exposed to α-MGH-1 antibody in both the 3 h and 24 h post 24 h dilution conditions, which was expected according to our peptide **1** data when peptides were free in solution. However, we have identified that APY took much longer to form and has not been observed after 3 h of incubation, only in the 24 h post 24 h dilution condition. Despite this, dark colored purple beads were still observed when beads were exposed to α-APY antibody, suggesting that the commercialized α-APY antibody is not specific to APY.

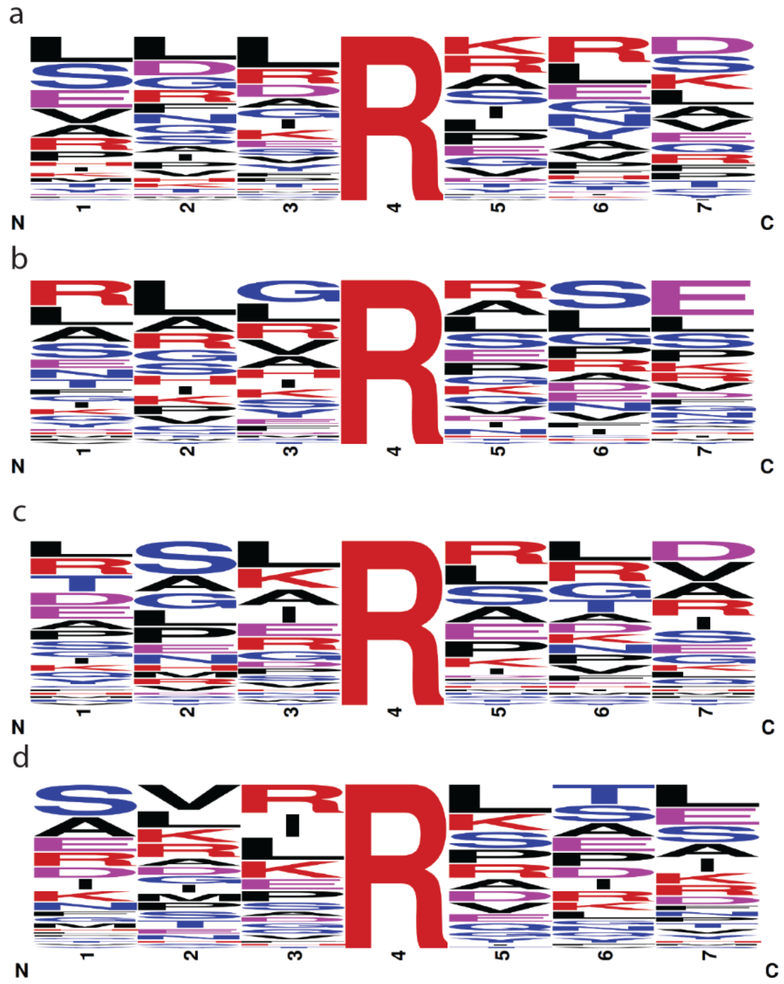

**Figure S21. Frequency Logos for All AGE-modified Peptides Observed in Proteomic Analysis.** Our prior work has shown that glycation is unlikely to have a strong consensus sequence, as it is largely driven by chemical microenvironments that include 3D protein structure. Nonetheless, it was also possible to evaluate sequence features of AGE-modified peptides. **(a)** Frequency consensus features for APY-modified peptides. **(b)** Frequency consensus features for MGH-modified peptides. **(c)** Frequency consensus features for CEA/MGH-DH-modified peptides. **(d)** Frequency consensus features for THP/other [M+144] AGE-modified peptides.

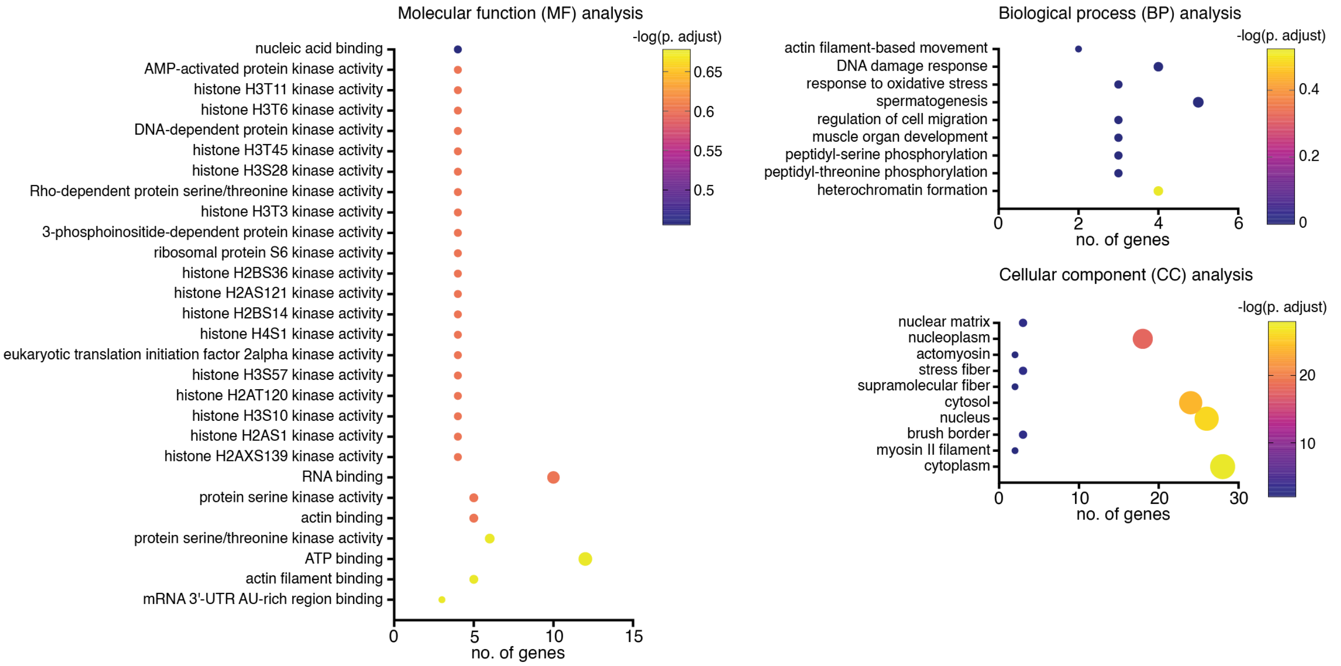


**Figure S22. DAVID GO Analysis Annotation Charts for APY Hits.** Each APY-modified peptide identified from our proteomic analysis was associated with one or more protein accession numbers (Uniprot). A list of all APY-containing accession numbers was input into DAVID for Gene Ontology (GO) analysis, and false discovery rate (FDR) was set to 5%. All GO terms were included in this figure. Multiple protein kinase activity terms in molecular function (MF) annotation and phosphorylation terms in biological process (BP) annotation appeared to be unique for APY hits. This observation is consistent with our *in vitro* peptide data, suggesting a connection between phosphorylation and APY formation in a cellular context.

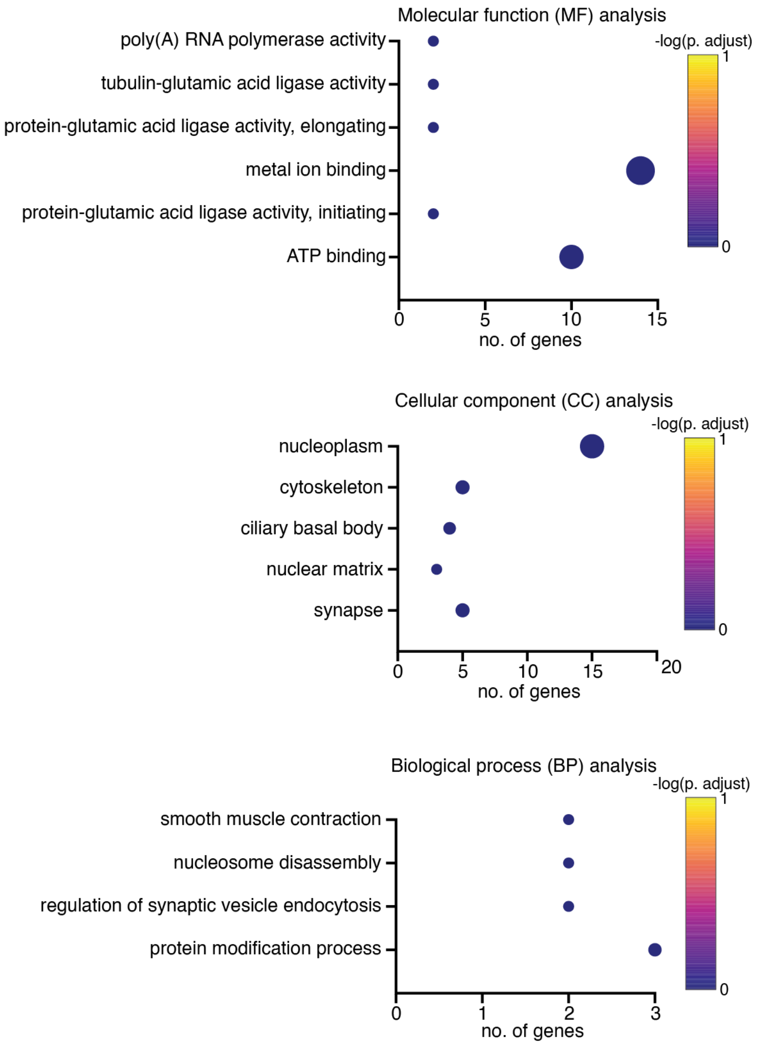


**Figure S23. DAVID GO Analysis Annotation Charts for MGH isomers Hits.** Each MGH-modified peptide identified from our proteomic analysis was associated with one or more protein accession numbers (Uniprot). A list of all MGH-containing accession numbers was input into DAVID for Gene Ontology (GO) analysis, and false discovery rate (FDR) was set to 5%. All GO terms were included in this figure.

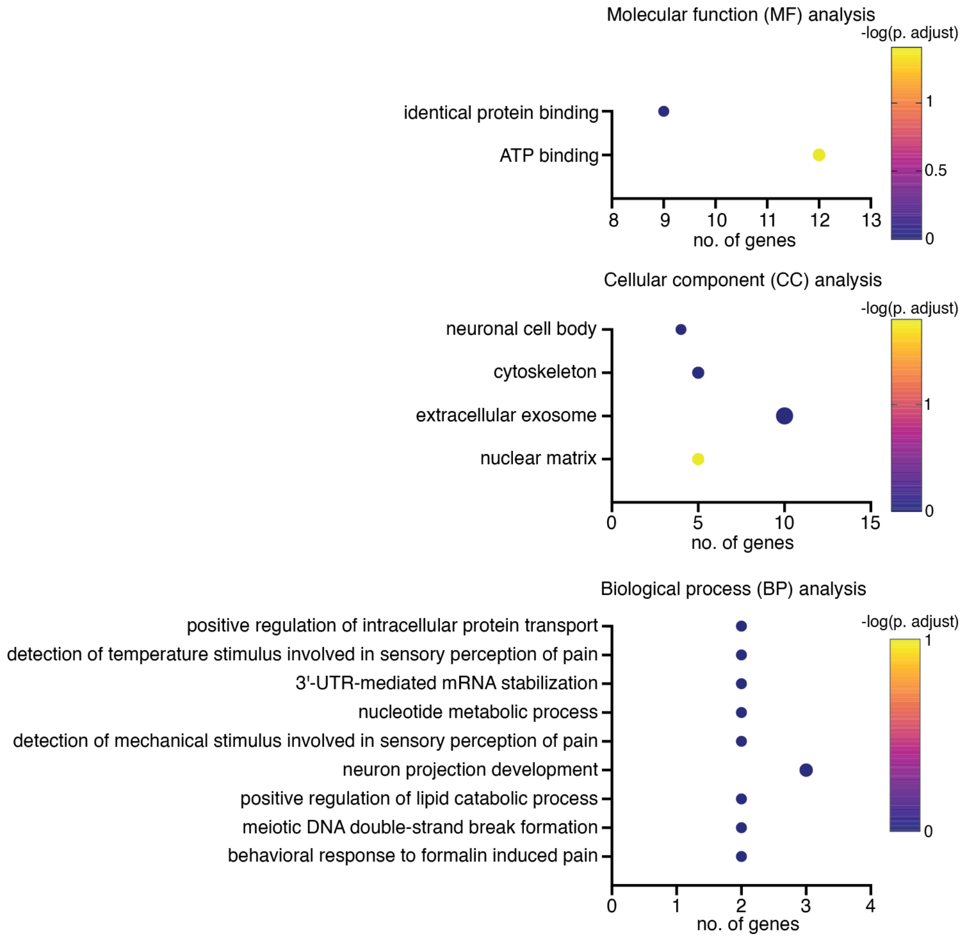


**Figure S24. DAVID GO Analysis Annotation Charts for CEA/MGH-DH Hits.** Although the dynamic modification for [M+72] adducts was set to include both Arg and Lys modifications, we only observed [M+72] adducts on Arg, corresponding to CEA and/or MGH-DH. Each CEA/MGH-DH-modified peptide identified from our proteomic analysis was associated with one or more protein accession numbers (Uniprot). A list of all CEA/MGH-DH-containing accession numbers was input into DAVID for Gene Ontology (GO) analysis, and false discovery rate (FDR) was set to 5%. All GO terms were included in this figure.

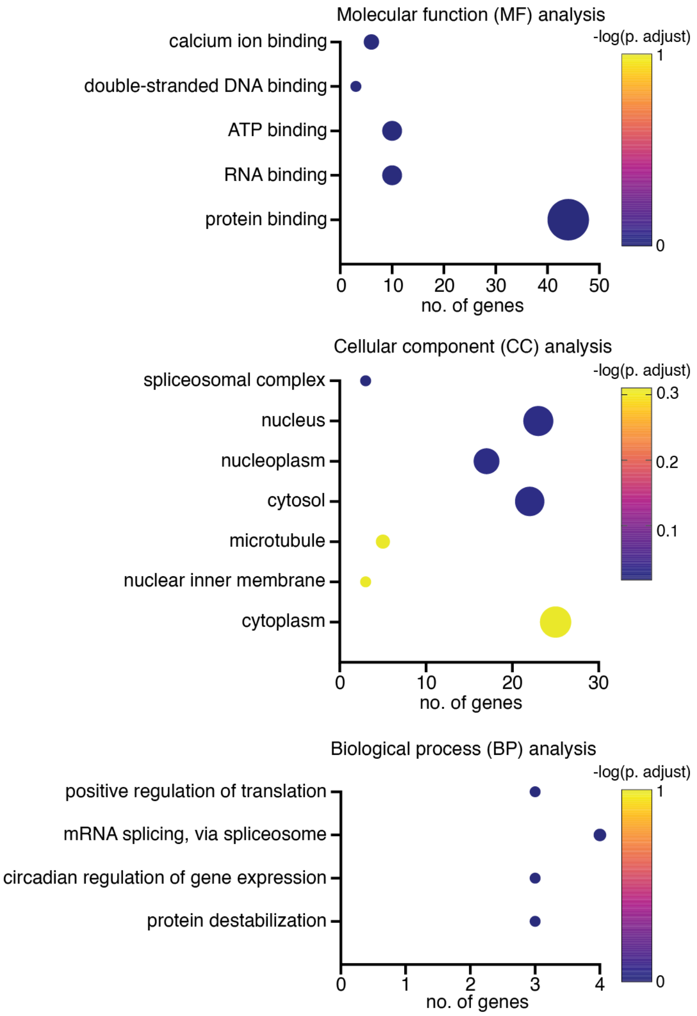


**Figure S25. DAVID GO Analysis Annotation Charts for [M+144] Hits.** Each THP/other [M+144] AGE-modified peptide identified from our proteomic analysis was associated with one or more protein accession numbers (Uniprot). A list of all [M+144]-containing accession numbers was input into DAVID for Gene Ontology (GO) analysis, and false discovery rate (FDR) was set to 5%. All GO terms were included in this figure.

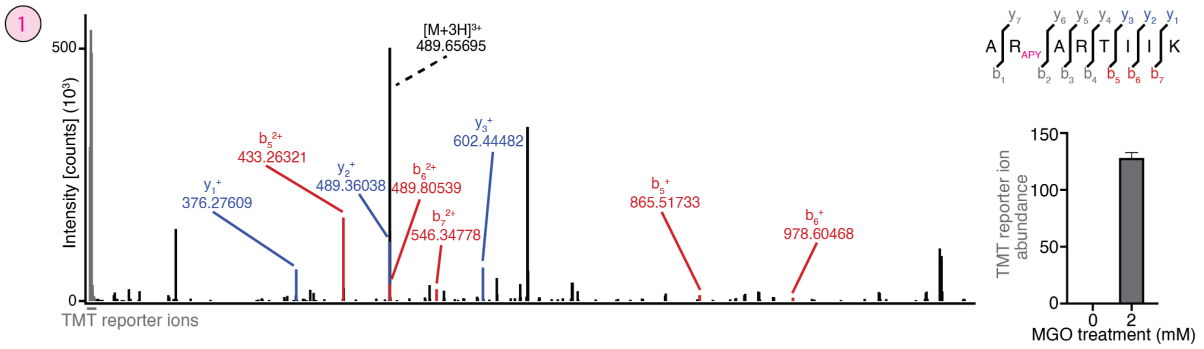


**Figure S26. b/y Ion Spectrum of APY-containing Unique Peptide ARARTIIK.**

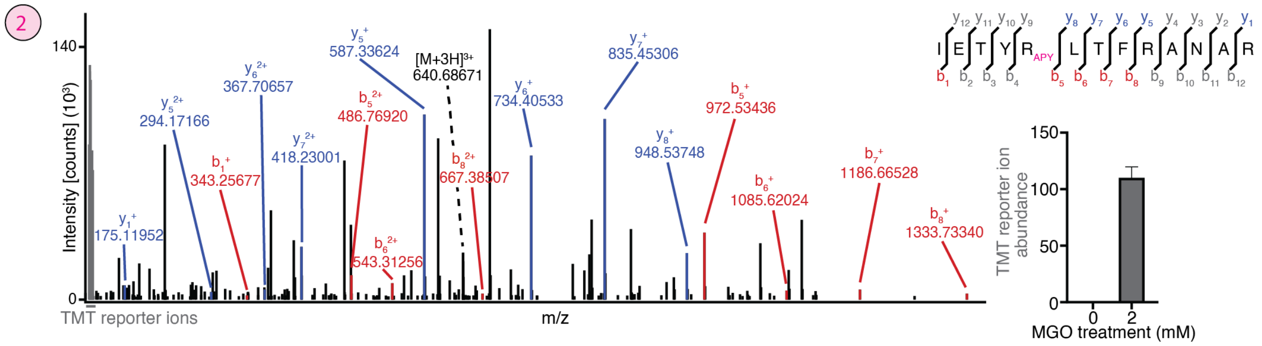


**Figure S27. b/y Ion Spectrum of APY-containing Unique Peptide IETYRLTFRANAR.**

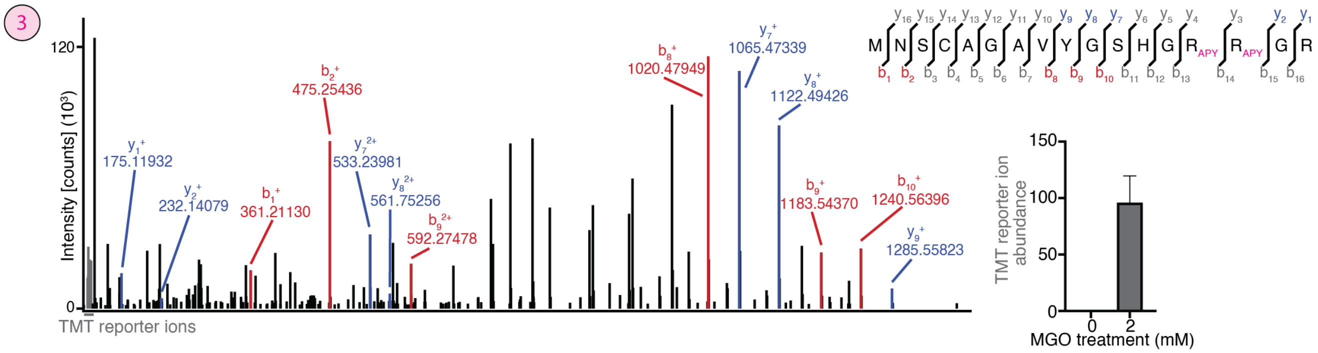


**Figure S28. b/y Ion Spectrum of APY-containing Unique Peptide MNSCAGAVYGSHGRRGR.**

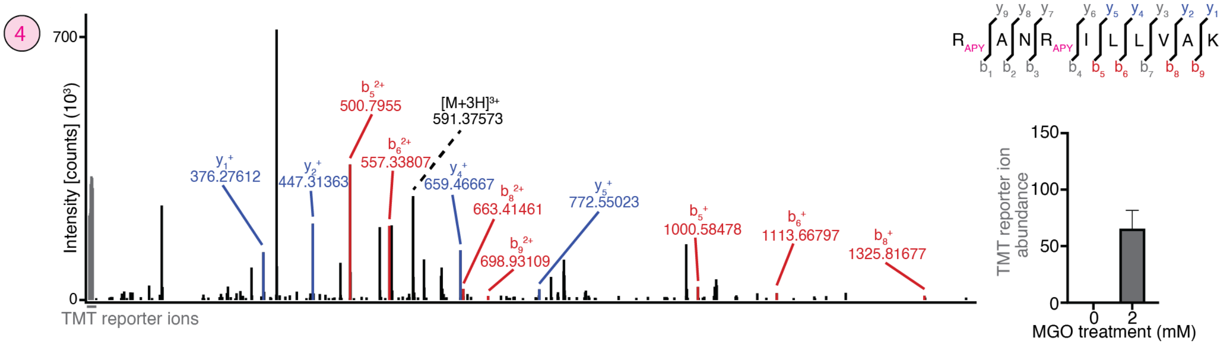


**Figure S29. b/y Ion Spectrum of APY-containing Unique Peptide RANRILLVAK.**

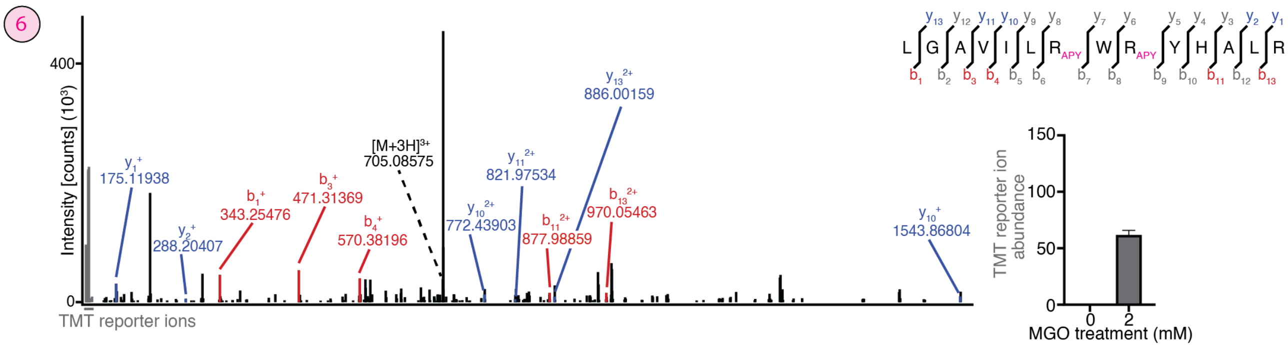


**Figure S30. b/y Ion Spectrum of APY-containing Unique Peptide LGAVILRWRYHALR.**

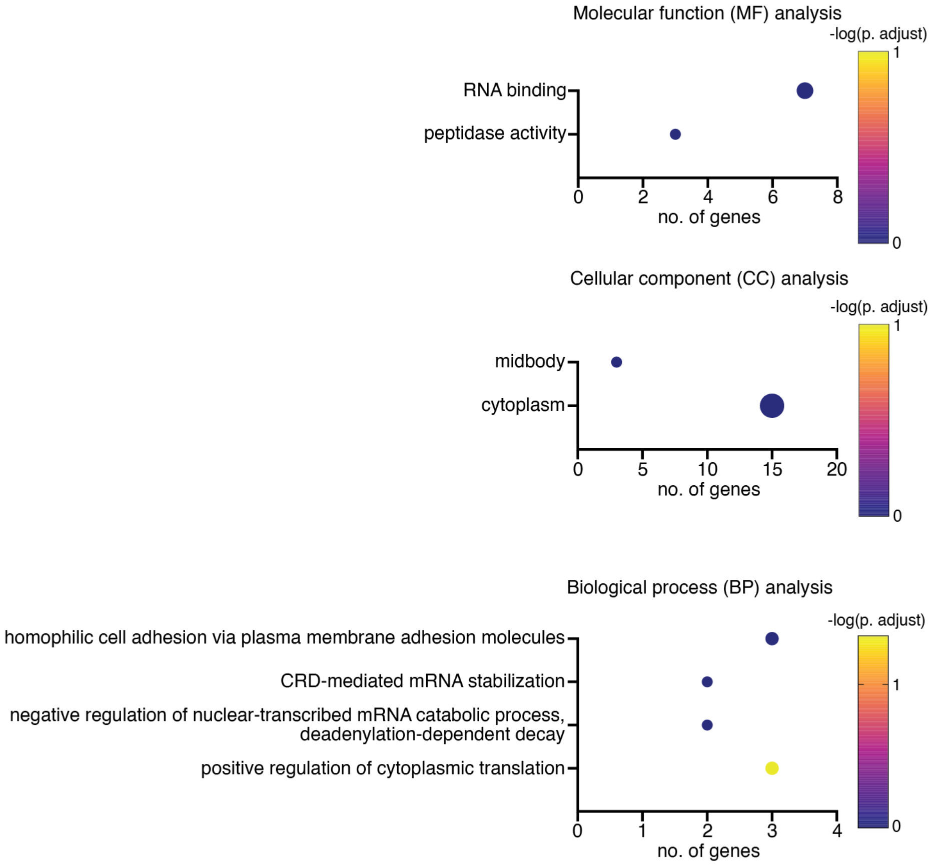


**Figure S31. DAVID GO Analysis Annotation Charts for Acetylation Hits.** Each Arg-acetylated peptide identified from our proteomic analysis was associated with one or more protein accession numbers (Uniprot). A list of all acetylation-containing accession numbers was input into DAVID for Gene Ontology (GO) analysis, and false discovery rate (FDR) was set to 5%. All GO terms were included in this figure.

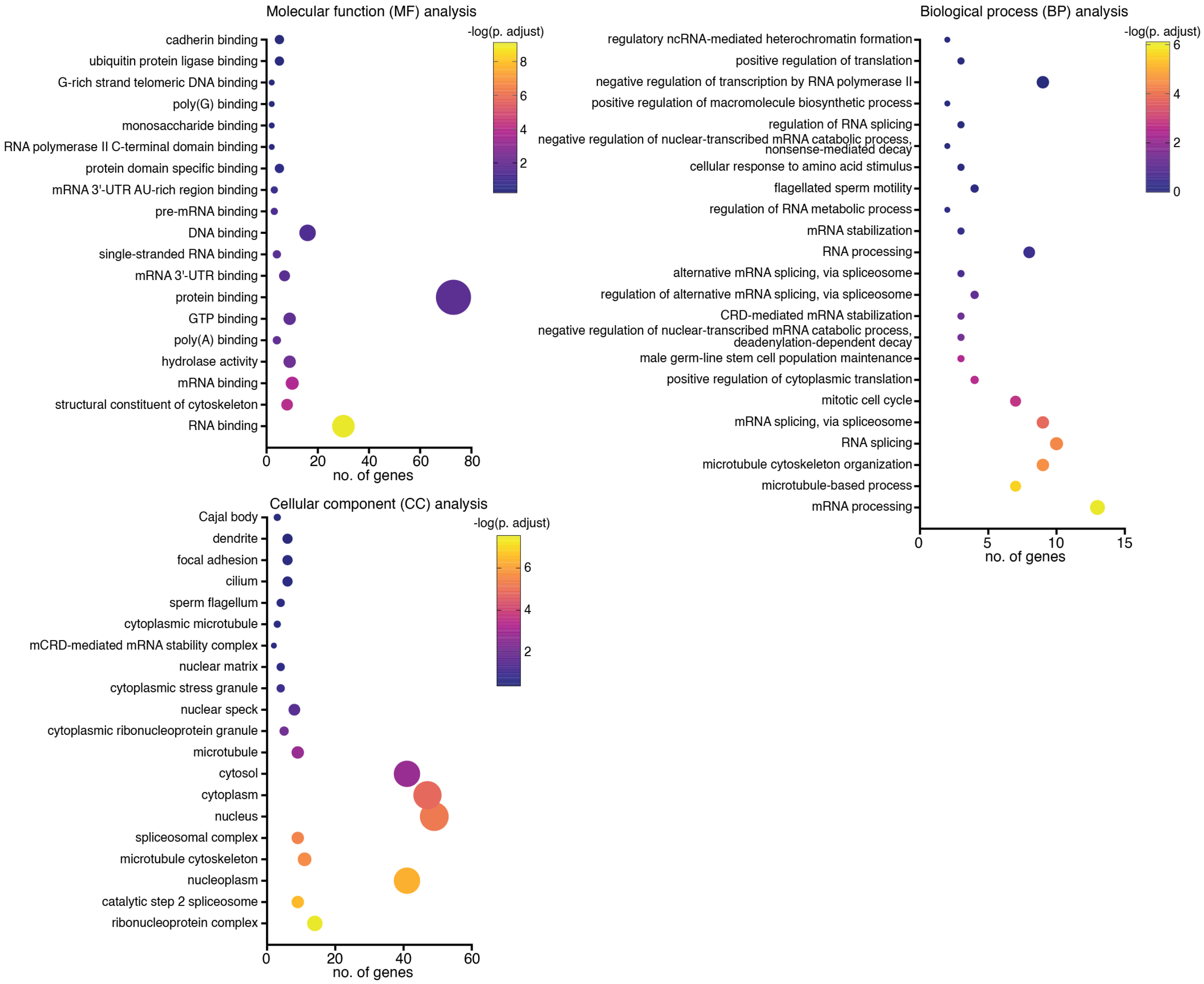


**Figure S32. DAVID GO Analysis Annotation Charts for Dimethylation Hits.** Each Arg-dimethylated peptide identified from our proteomic analysis was associated with one or more protein accession numbers (Uniprot). A list of all dimethylation-containing accession numbers was input into DAVID for Gene Ontology (GO) analysis, and false discovery rate (FDR) was set to 5%. All GO terms were included in this figure.

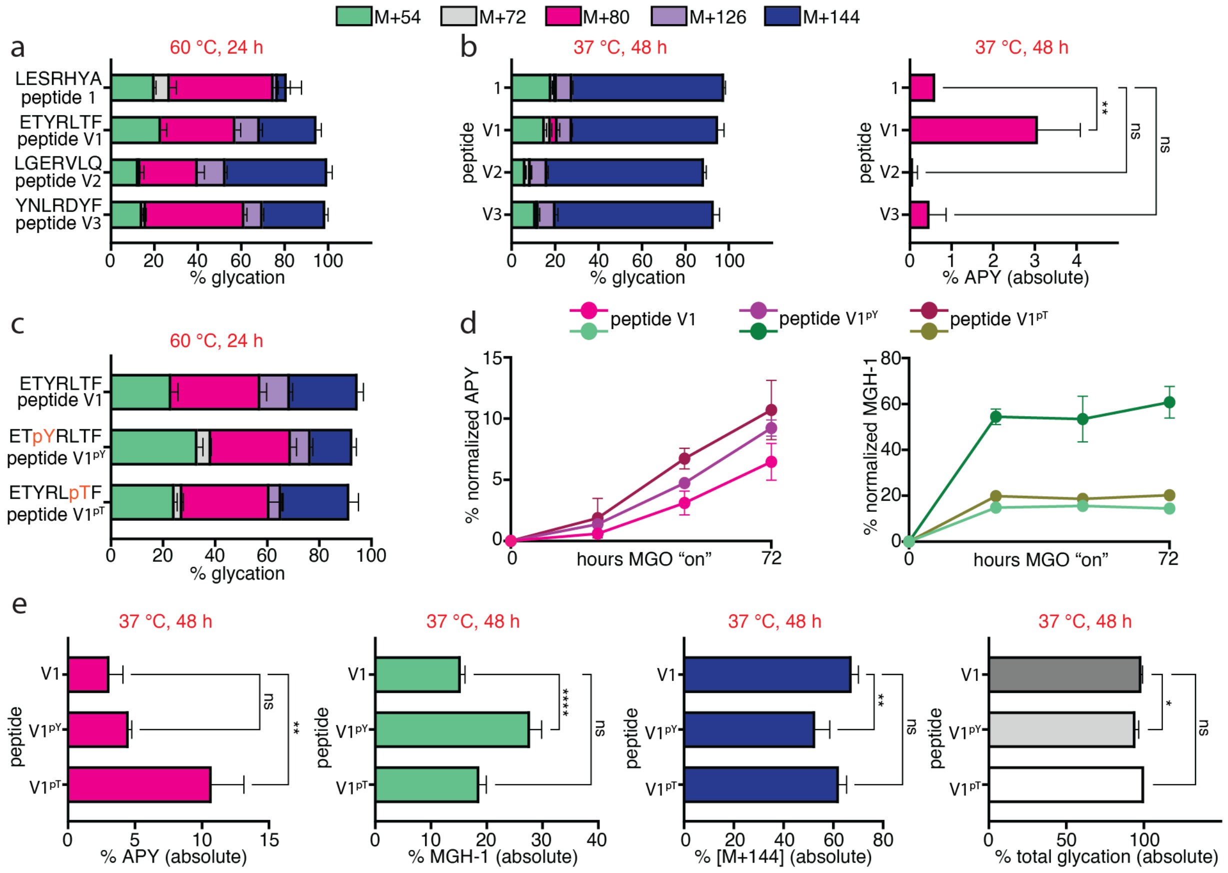


**Figure S33. Proteomic Analysis Validation with in vitro Peptide Data.** In order to validate our proteomic analysis, we synthesized three peptides found to be APY-modified in our dataset: Ac-ETYRLTF from MAP4 (hit 2; peptide **V1**), Ac-LGERVLQ from ELAVL3 (hit 7; peptide **V2**), and Ac-YNLRDYF from HNRNPA3, a non-hit yet high-abundant sequence (peptide **V3**). (**a**) Distributions of AGEs at 60 °C at 24 h for peptides **1**, **V1**, **V2**, and **V3** (n = 3). All three peptides were confirmed to form APY. Expectedly, peptide **V2**, without a nearby Tyr, exhibited the least APY formation out of the three peptides, while peptide **V3**, with two Tyr residues, exhibited the most APY formation. (**b**) Distributions of AGEs at 37 °C at 48 h (right stacked bar graph) for peptides **1**, **V1**, **V2**, and **V3**, and absolute APY levels (%) observed under the same conditions (left bar graph) (n = 3 for all reactions). Under these conditions, peptide **V1**, our high-confidence hit, showed the most APY formation, followed by peptides **1** and **V3**, and expectedly, peptide **V2** formed the least APY. Ordinary one-way ANOVA was used to determine if each variant yielded statistically significant differences in % absolute APY compared to peptide **1**. $p$<0.01(**). (**c**) Distributions of AGEs at 60 °C at 24 h for peptides **V1**, **V1$^{pY}$**, and **V1$^{pT}$** (n = 3). In order to further investigate the crosstalk between glycation and phosphorylation, we synthesized two variants of peptide **V1** that incorporated either pTyr or pThr (Ac-ETpYRLTF, peptide **V1$^{pY}$**, and ETYRLpTF, peptide **V1$^{pT}$**). All three peptides exhibited similar levels of overall glycation and APY formation, despite the introduction of extra negative charges. (**d**) Time course data (% normalized APY, left; % normalized MGH-1, right) at 37 °C for peptides **V1**, **V1$^{pY}$**, and **V1$^{pT}$** taken over 24 h, 48 h, and 72 h (n = 3). Peptide **V1$^{pT}$** was the fastest to form APY, followed by peptide **V1$^{pY}$**, and then peptide **V1**. (**e**) Ordinary one-way ANOVA was used to determine if each variant yielded statistically significant differences in % absolute APY (left), % absolute MGH-1 (middle left), % absolute [M+144] (middle right), and % absolute total glycation (right) of **V1$^{pY}$** and **V1$^{pT}$** compared to peptide **V1** (n = 3). $p$<0.05(*), $p$<0.01(**), $p$<0.0001(****). Absolute APY levels in peptide **V1$^{pT}$** were three-fold higher than those in peptide **V1**, and both phosphorylated variants exhibited an increase in MGH-1 formation, indicating that phosphorylation can also bias the formation of certain AGEs.
